# Axon Regeneration and Functional Recovery after Spinal Cord Injury is Enhanced by Allele-Specific ApoE Neuronal Action through LRP8

**DOI:** 10.1101/2025.10.10.681747

**Authors:** Ramakrishnan Kannan, Xingxing Wang, LaShae Nicholson, Nancy Z. Lin, Elisa M. Howard, Atrayee Basu, Ines Ingabire, Yuichi Sekine, Stephen M. Strittmatter

## Abstract

Adult CNS trauma frequently causes neuronal disconnection and persistent deficits due to failed axon regeneration. While model system screening has identified multiple candidate neural repair pathways, ApoE–LRP8 signaling is unique in being implicated clinically. Here, we show that cortical axon regeneration requires LRP8 and is modified by *APOE* variants. ApoE2-expressing mice show reparative corticospinal and raphespinal axon growth with greater motor function than controls after spinal cord injury. Distinct from ApoE in other settings, there is no change in inflammation or scarring. After axotomy, ApoE exerts allele-specific effects on LRP8 localization and signaling in cortical neurons. *APOE* alleles regulate synaptic organization gene expression by cortical neurons after injury, with little effect on glial gene expression. AAV-mediated overexpression of ApoE2 in mice after spinal trauma increases locomotor recovery and reparative axon growth. Thus, ApoE–LRP8 signaling for axon regrowth following CNS trauma provides a potential therapeutic intervention site.

## INTRODUCTION

Many acute injuries to the central nervous system interrupt axonal connectivity. This is most dramatic in traumatic spinal cord injury (SCI) where very few neurons are lost but communication between the brain and distal spinal cord are nil after trauma, and neurological deficits can be pronounced(*1*). Unfortunately, repair of connectivity through axonal regeneration is extremely limited in adult mammals so recovery is typically poor(*2*). While overcoming limitations to reparative axon growth provides a possible avenue towards clinical improvement, the 300,000 individuals in the United States with persistent neurologic deficits after SCI have no therapeutic option today(*3*).

A range of non-clinical experiments have sought to define molecular restraints upon axonal growth and plasticity after injury(*4, 5*). Potential molecular approaches to overcome the brakes on neural repair have not yet entered clinical practice, though several promising approaches are now being tested in clinical trials(*6, 7*). This includes blockade of myelin-derived inhibitors and interruption of chondroitin sulfate proteoglycan action. Training and electrical stimulation have been used to coax neurons into a more plastic state to support functional recovery and have achieved early clinical promise(*8, 9*). Despite these advancements there remains no therapeutic intervention to improve recovery after traumatic spinal cord injury.

We conducted a genome wide loss of function screen in cortical neurons to identify those genes modifying the extent of axonal regeneration *in vitro*(*10*). Over 500 genes with significant effects were identified and dozens were validated with an *in vivo* optic nerve regeneration screen(*11*). Rab27B(*10*), rabphilin-3A(*12*) and interleukin-33(*11*) were further confirmed using gene knockout mice in spinal cord injury or optic nerve crush experiments. In addition, specific experiments have shown that PTEN(*13*), SOCS3(*14*), KLF4(*15*), neurotrophins, and several transcription factors titrate the regenerative capacity of CNS axons(*16–18*). Amongst these numerous candidates for enhancing neural repair, evidence for contribution to clinical recovery via genetic association would be extremely helpful to validate and assess relevance.

Genome wide association studies to seek variants that might modulate the neural repair process have not been reported for SCI, but studies of traumatic brain injury (TBI) have been completed. The ApoE4 variant reached genome-wide significance for TBI outcome(*19*), and has been linked to SCI recovery clinically and experimentally(*20–23*). In our screen(*10*), loss of the ApoE receptor LRP8 suppressed axon regeneration. Here, we examined whether an ApoE–LRP8 signaling axis might regulate the extent of neural repair after spinal cord injury. We report that ApoE has an allele-specific effect on an LRP8-dependent axonal regeneration pathway. ApoE2 knock-in mice show selective changes of neuronal gene expression, enhanced axon regeneration and greater functional recovery after traumatic spinal cord injury. Furthermore, increasing ApoE2 after trauma enhances neural repair and recovery. Notably, the neuronal selectivity of ApoE–LRP8 action in SCI recovery differs from ApoE action on protein clearance, transport, cell metabolism and inflammation in neurodegenerative studies, indicating the diverse action of ApoE in multiple neurological conditions.

## RESULTS

### ApoE ligand and neuronal LRP8 receptor are indispensable for axon growth

Our previous genome-wide shRNA screen identified dozens of genes for which loss of function modified the success of axon regeneration *in vitro* as well as *in vivo* after optic nerve crush and spinal cord trauma(*10–12*). Within the Low-density lipoprotein Receptor-related Protein (LRP) family, suppression of *Lrp6* and *Lrp8* expression significantly reduced axon regeneration (Fig. S1A). LRP8 is known to be enriched in neurons and to modulate brain development and adult synaptic plasticity(*24–27*). Consistent with the shRNA data, deletion of mouse *Lrp8* (Fig. S1B) inhibits axon regeneration from cortical neurons relative to wild-type in a gene dosage-dependent manner (Fig. 1A and B).

**Fig. 1.**
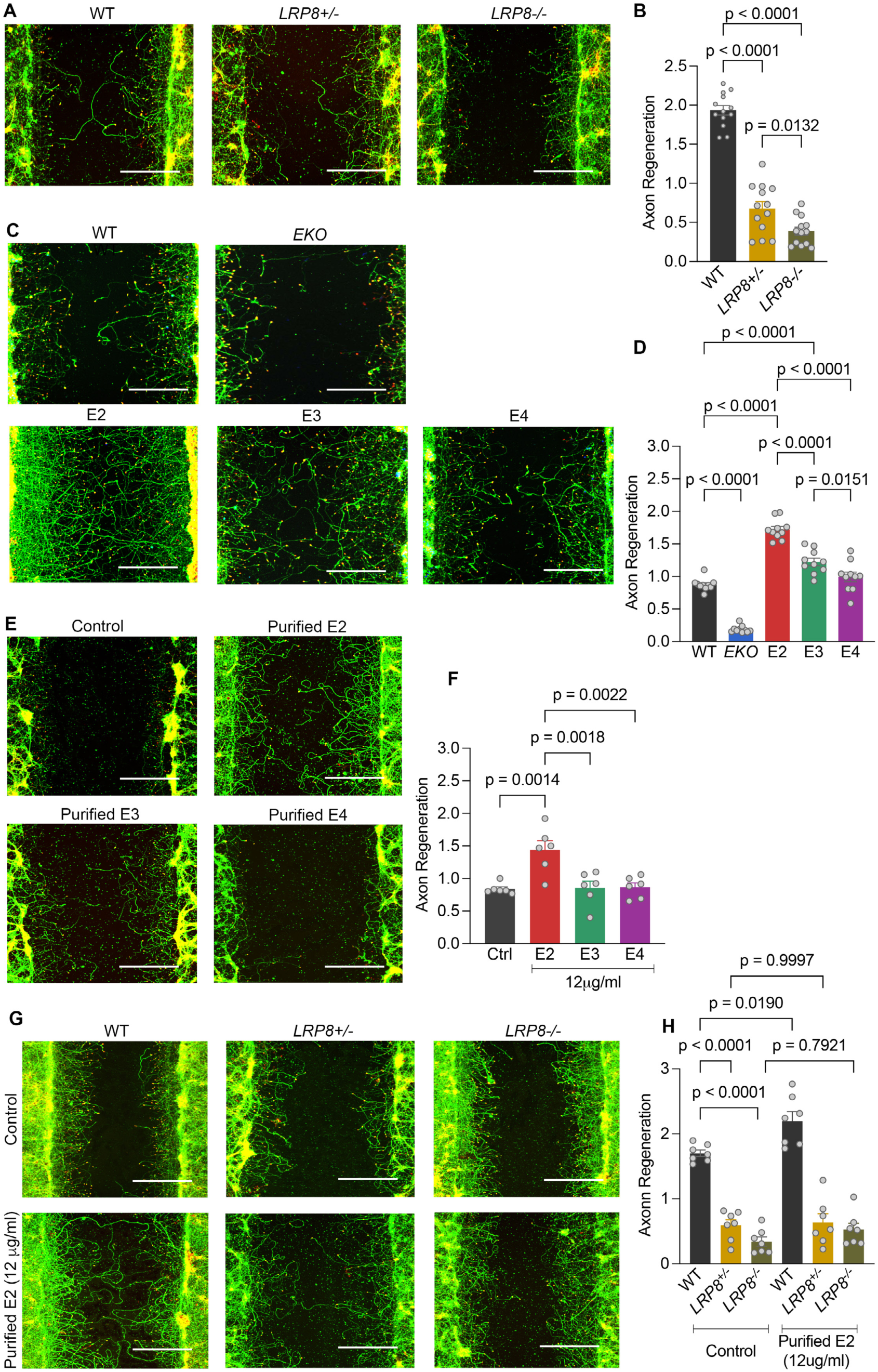
E2 promotes axon regeneration through neuronal LRP8 receptor. Axotomized primary cortical neuron cultures stained with ßIII-tubulin (green) and phalloidin (red) at 15 days after plating and 7 days after axotomy. A portion of regeneration zone is shown here. For each genotype, datapoints are an average of 60 wells from independent biological replicates. Regeneration scores are normalized to WT. Scale bars, 200 µm. (A) Photomicrographs of axon regeneration from WT, *LRP8^+/-^* and *LRP8^-/-^* cortical neuron cultures. (B) Axon regeneration index for (A) from 13 mice for each group. (C) Representative immunofluorescence micrographs of axon regeneration from WT, EKO, E2, E3 and E4 cortical neuron cultures. (D) Axon regeneration index for (C) from 10 mice for each group. (E) Photomicrographs of axon regeneration from EKO cortical neurons treated with purified ApoE lipoprotein particles (12 μg/ml) from astrocytic conditioned media (ACM). (F) Axon regeneration index for (E) from 6 mice for each group. (G) Photomicrographs of axon regeneration from WT, *LRP8^-/+^* and *LRP8^-/-^* cortical neuron cultures treated with purified E2 lipoparticles with equal amount of PBS as control. (H) Axon regeneration index for (E) from 7 mice for each group. Data presented as mean ± SEM. One-way ANOVA with Tukey’s multiple comparisons for (B): F = 145.2, dF = 2; (D): F = 127.4, dF = 4; (F): F = 9.3, dF = 3; (H): F = 54.5, dF = 5.

ApoE and Reelin are known physiological ligands for LRP8 receptor in the brain(*28*), where ApoE is a major lipid and cholesterol transporter. However, the role of ApoE isoforms in CNS regeneration has yet to be fully elucidated. We considered whether ApoE isoforms have varying effects on regenerative axon growth via LRP8. Axon regeneration after mechanical trauma was assessed for cortical neurons cultured from mice homozygous for knock-in (KI) of the three common human *APOE* alleles (E2, E3 or E4) or from *ApoE* null mice (EKO) or from wild-type mice. The E2 cortical neuron cultures display a significant increase in axon regeneration, while the EKO neurons exhibit reduced regeneration (Fig. 1C and D). Taken together, these data show that ApoE and its receptor LRP8 are crucial for axon regeneration with expression of ApoE2 stimulating growth.

### Secreted lipidated E2 modulates LRP8 signaling for reparative axon growth

Astrocytes are known to be the primary source of secreted lipidated ApoE in the brain(*29*). We sought to evaluate the role of astrocyte-secreted ApoE in axon regeneration. Cortical neuron cultures utilized for axon regeneration assays *in vitro* contain nearly 90% neurons, with astrocytes being the most prevalent glial cell(*10*). As a first step, we assessed cell type specific differences in expression from human *APOE* knock-in alleles by immunoblotting tissue homogenates as well as cellular and secreted fractions of separately cultured cortical neurons and astrocytes. In cortical tissue, the levels of ApoE were uniform across the three different *APOE* knock-in mice (Fig. S1C). The expression and secretion of E2 was similar between the different fractions (Fig. S1D). The level of E4 was reduced in cellular and secreted neuronal extracts as compared to E3 (Fig. S1E), but was comparable for astrocyte cultures (Fig. S1F).

To test the role of secreted ApoE in axon regeneration, we used astrocyte-conditioned medium from the three humanized ApoE-KI models together with EKO neuron cultures, thereby eliminating any influence of neuronal ApoE or wild-type mouse ApoE. Cultures of EKO astrocytes yielded control ApoE-free medium. Astrocyte-conditioned medium extracts from E2 and E3 facilitated regeneration of EKO neurons more effectively than does E4 medium (Fig. S1G and S1H). To exclude the possibility of differential growth factor levels in astrocyte-conditioned medium and to establish the primary role of lipidated secreted human ApoE in regeneration, we immuno-purified ApoE from astrocyte-conditioned medium. Native gel electrophoresis of purified fractions revealed predominant association of ApoE isoforms with high molecular weight lipid-rich particles (Fig. S1I).

Purified E2 lipoprotein particles substantially increased axon regeneration of EKO neurons compared to E3 and E4 (Fig. 1E and 1F). To determine whether E2 lipoparticles signal through neuronal LRP8 receptor, we conducted an *in vitro* epistasis experiment with wild-type and LRP8 null neurons. Secreted E2 significantly enhanced axon regeneration of wild-type but not *Lrp8^-/-^* and *Lrp8^-/+^* cortical neurons indicating that E2 modulates axon regeneration through neuronal Lrp8 signaling (Fig. 1G and H).

### E2 mice show enhanced CNS regeneration with improved functional recovery from SCI

We hypothesized that the effect of ApoE variants on recovery from clinical TBI and SCI relates to titration of reparative axonal growth through neuronal LRP8, and sought to test this for mouse SCI. Before investigating anatomical and behavioral SCI recovery, we confirmed that the development of two relevant descending spinal tracts was normal without injury. Both corticospinal (CST) and raphespinal projections show normal anatomy in the three ApoE-KI strains and in ApoE null mice (Fig. S2A and S2B). Therefore, to assess the role of ApoE alleles in CNS repair and functional recovery *in vivo*, we performed thoracic spinal cord dorsal over-hemisection in adult ApoE-KI mice (Fig. 2A). By Basso Mouse Score (BMS) locomotor evaluation in the open field, all experimental groups had normal hindlimb function before injury, but essentially complete paralysis 3 days after injury. We tracked locomotor ability of injured mice until 63 days post-SCI (Fig. 2B). From 21 days onwards, recovery of the E2 group was significantly greater than for WT or E4 mice (Fig. 2B). Nearly all E2 mice achieved extensive ankle movement and hindlimb support while most WT and E4 mice exhibited limited ankle movement and absent hindlimb weight support. Locomotor recovery in E3 mice was indistinguishable from WT mice, and the EKO mice had less restoration of hindlimb function after injury (Fig. S2C). In addition, we assessed interlimb coordination during locomotion through a linear track by CatWalk analysis at 8-wks post-SCI (Fig. 2C). Consistently, E2 mice displayed better limb coordination and balance with paw support compared to WT and E4 mice (Fig. 2D). The automated analysis of paw prints detected hindlimb steps for more than 75% of forelimb steps in E2 mice but fewer than 5% of forelimb steps of WT and E4 mice. Thus, ApoE expression has a prominent role in SCI recovery with ApoE2 being most beneficial for locomotor outcome, and with ApoE3, ApoE4 and mouse ApoE each supporting more improvement than the ApoE null state.

**Fig. 2.**
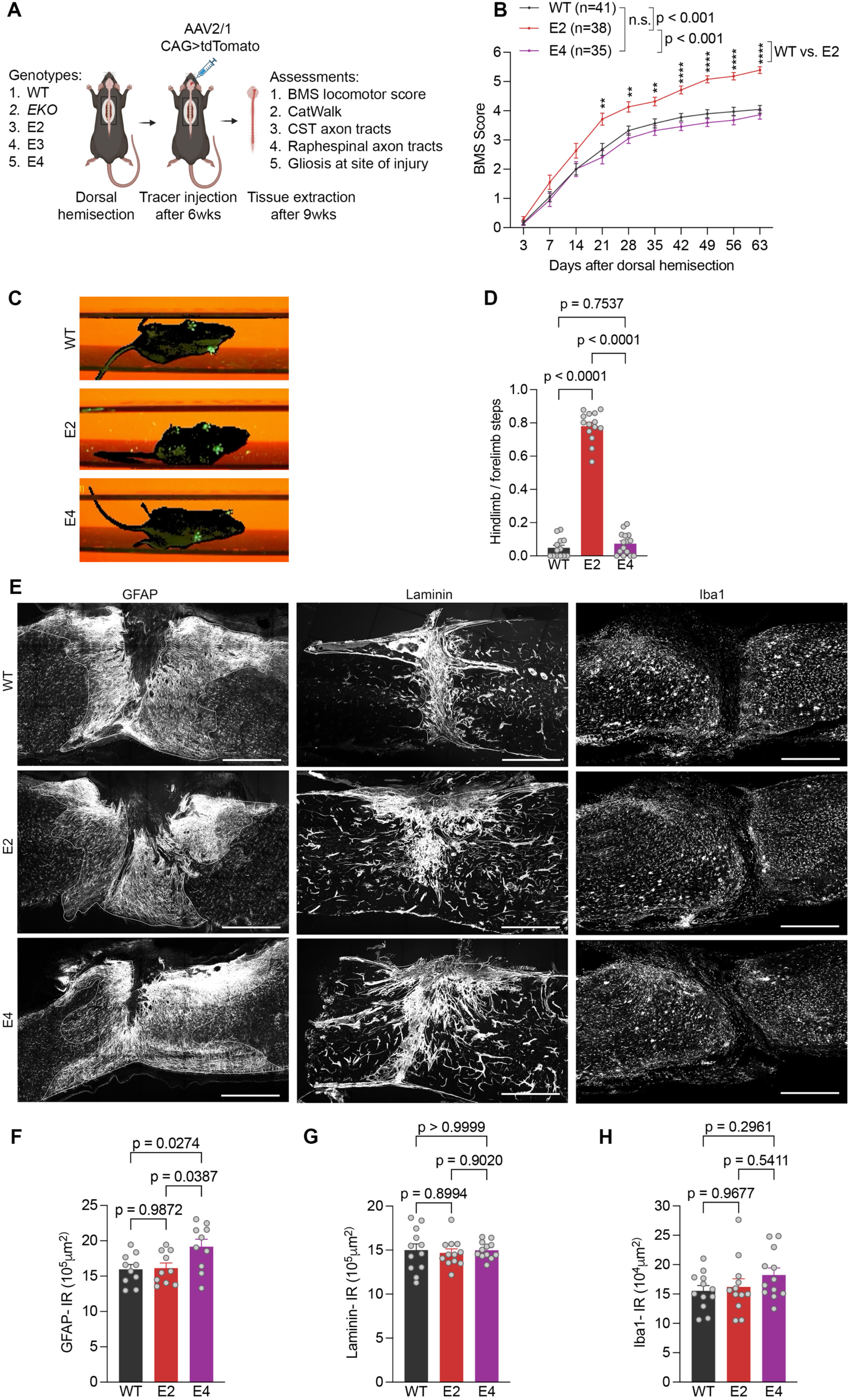
Enhanced locomotor recovery in E2 mice after dorsal thoracic over-hemisection. (A) Experimental overview to evaluate functional and anatomical features of motor recovery in SCI mice with different ApoE-KI alleles. (B) BMS open-field locomotion assessment for WT (*n* = 42), E2 (*n* = 38) and E4 (*n* = 35) SCI mice. Performance was scored for each animal every consecutive week. Data as mean ± SEM and *p* values calculated by repeated measure multi-point ANOVA across time series followed by post hoc Tukey’s multiple comparisons test between genotypes at indicated time points. ****p<0.0001, **p<0.01 (C) Single frame of CatWalk video showing fore and hindlimb paw footprints in green for WT, E2 and E4 mice 9 weeks after SCI. (D) Limb coordination index of WT (*n* = 13), E2 (*n* = 14) and E4 (*n* = 15) mice, 9 weeks after SCI. (E) Photomicrographs of sagittal spinal cord sections at thoracic SCI epicenter. Inflammation was assessed by quantitating GFAP, Laminin and Iba1 immunoreactive area at the injury site for *n* = 10 WT, E2 and E4 mice. Scale bar, 500 µm. (F) Quantification of GFAP immunoreactive area at injury site. (G) Quantification of Laminin immunoreactive area at injury site. (H) Quantification of Iba1 immunoreactive area at injury site. Data shown as mean ± SEM and *p* values calculated by one-way ANOVA with Tukey’s multiple comparisons test for (D): F = 455.1, dF = 2; (F): F = 4.77, dF = 2; (G): F = 0.13, dF = 2; (H): F = 1.45, dF = 2.

After SCI, reactive astrocytes, microglia, peripheral immune cells and fibroblasts invade the SCI lesion core and alter tissue damage during the acute and subacute phases. Therefore, ApoE-regulated locomotor recovery might be attributable to either differential neuroprotection or reparative axonal growth, though the late behavioral recovery favors a neural repair mechanism rather than neuroprotection. As assessed by GFAP-positive astrocytosis, fibroblast-derived laminin deposition or Iba1-positive microglial presence, there was no difference between the spinal cord lesion sites of E2, E4 and WT mice (Fig. 2E-2H). This further supports the hypothesis that axonal regrowth rather than neuroprotection accounts for improved BMS after SCI.

To evaluate axonal growth as a function of ApoE, we assessed descending CST projections from motor cortex and 5-HT tracts from hindbrain raphe neurons, both of which are known to contribute to functional performance after SCI. To trace the CST unilaterally, we injected AAV-mCherry into the sensorimotor cortex 9 weeks after SCI and collected tissue 4 weeks later at 13 weeks post-SCI. There was equally robust labelling of CST axons rostral to the injury site in parasagittal sections of the thoracic spinal cord and in transverse sections of the cervical cord for the E2, E4 and WT mice (Fig. 3A and 3B). At 400 µm caudal to the injured site there were rare fibers in WT mice, numbering 0.2% of the cervical count (Axon Index of 0.002) and no fibers were detected in E4 mice (Fig. 3A and 3C). There were three times more CST fibers caudal to the injury in the E2 cohort as compared to the WT mice. Examples from multiple additional mice are shown in Fig. S3A. Caudal CST fibers in EKO and E3 mice at 13 weeks after SCI were as rare as in WT mice (Fig. S3C).

**Fig. 3.**
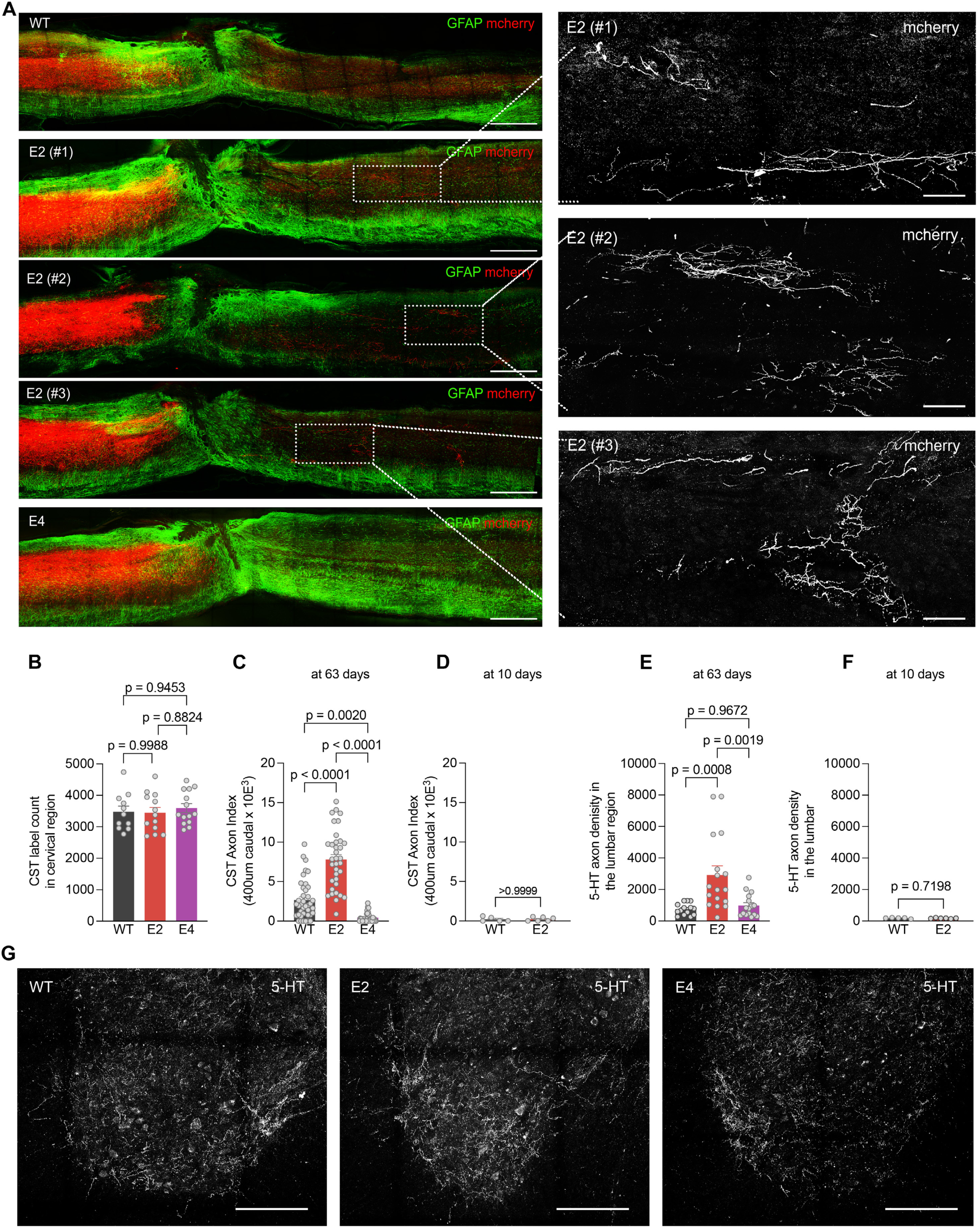
Enhanced regeneration of corticospinal and raphespinal axon tracts in E2 mice after SCI. (A) Sagittal photomicrographs of spinal cord spanning the thoracic dorsal over-hemisection lesion at 63 days after injury. AAV-mCherry anatomical tracer used to label regenerating CST axons (red), and GFAP immunohistology for reactive astrocytes (green). Dorsal is up and rostral is left. Scale bar, 500 µm. White outlined boxed areas in each image are captured at high resolution to visualize regenerating CST fibers caudal to lesion for red channel only. Scale bar, 100 µm. (B) Quantification of CST labelling efficiency using anatomical tracer at the cervical region of injured spinal cord for WT (*n* = 11), E2 (*n* = 13) and E4 (*n* = 14). (C) Quantification of CST axons based on anatomical tracer in lumbar spinal cord caudal to lesion 63 days after SCI with tracer injection at day 42. CST fibers were assessed 400 µm caudal to the lesion site in WT (*n* = 39), E2 (*n* = 35), E4 (*n* = 31). (D) Quantification of spared CST axons at 10 days after SCI, traced by biotin-dextran-amine tracer injection at post-injury day 1, in lumbar spinal cord 400 μm caudal to lesion in WT (*n* = 5) and E2 (*n* = 6). (E) Quantification of serotonergic (5-HT+ve) axon length at 63 days after SCI in the ventral horn of lumbar spinal cord for WT (*n* = 14), E2 (*n* = 17), E4 (*n* =16). (F) Quantification of spared serotonergic (5-HT+ve) axon length at 10 days after SCI in the ventral horn of lumbar spinal cord for WT (*n* = 5) and E2 (*n* = 6). (G) Transverse section photomicrograph of ventral horn lumbar spinal cord at 63 days after injury stained with anti-5-HT antibody. Scale bar, 100 µm. Data shown as mean ± SEM. *p* values calculated by one-way ANOVA with Sidak’s multiple comparisons test for (B): F = 0.24, dF = 2; (C): F = 68.48, dF = 2; (E): F = 10.01, dF = 2 and by two-tailed unpaired t-test for (D): t = 0.09, df = 8; (F): t = 0.37, dF = 9.

We considered whether the observed E2 CST fibers grew after injury, or were initially spared from axotomy. In separate cohorts of mice from which tissue was collected 10 days after injury, CST tracing initiated with biotin-dextran-amine injection one day after injury showed robust labelling immediately rostral to the injury, but no fibers below the injury in WT and E2 mice (Fig. 3D). Coupled with the normal development of the CST in ApoE mutant mice (Fig. S2A), these data demonstrate that the caudal CST fibers of E2 mice derive from post-SCI axon growth either via long-distance axon regeneration or via regenerative sprouting from rare uninjured fibers.

We also visualized raphespinal fibers by anti-5HT staining in the ventral horn of the lumbar enlargement. The dorsal over hemisection injury interrupts many, but not all of these axons, and in WT mice there is a reduced density of 5HT axons in the lumbar ventral gray matter (Fig. 3E and 3G). The density of raphespinal fibers is increased more than 3-fold in E2 mice as compared to WT, despite the normal raphespinal development in uninjured E2 mice (Fig. S2B). The E4 genotype had no effect on 5HT innervation density in the caudal cord after SCI. Multiple additional examples are shown in Fig. S3B. Analysis of lumbar raphespinal fibers in mice sacrificed at 10 days post-SCI showed few 5HT axons in the lumbar cord with no difference in the between genotypes (Fig. 3F). These findings are consistent with selective enhancement of distal axon sprouting in the E2 cohort at 13 weeks post-injury. We conclude that ApoE alleles influence the extent of axon regrowth and functional recovery after CNS trauma. The E2 allele promotes reparative axon growth, while E4 allele inhibits axon extension.

### ApoE isoform effects of LRP8 trafficking and signaling in axon regrowth

Based on the cell culture axotomy data, we hypothesized that increased CST and raphespinal regrowth after injury was due to altered ApoE–LRP8 signaling in neurons subject to axotomy by SCI. We explored LRP8 subcellular localization as a marker of altered receptor signaling. The distribution of LRP8 in L5 neurons of M1 cortex from E2, E3 and E4 at 6 weeks after SCI was compared to naïve controls (Fig. 4A, 4B). While all genotypes tested had similar overall LRP8 levels in M1 cortex pre- and post-SCI, the effect of SCI was significantly different between ApoE genotypes. The E3 and E4 cortex showed a pronounced shift of LRP8 immunoreactivity from neuropil to neuronal cell soma after SCI, while an altered subcellular distribution was not detected in E2 samples. By contrast, the subcellular distribution of LRP5 and LRP6 receptors in the cerebral cortex was not altered after SCI or with different ApoE genotypes (Fig. S4A-S4D). These data demonstrate ApoE allele-specific regulation of LRP8 after SCI.

**Fig. 4.**
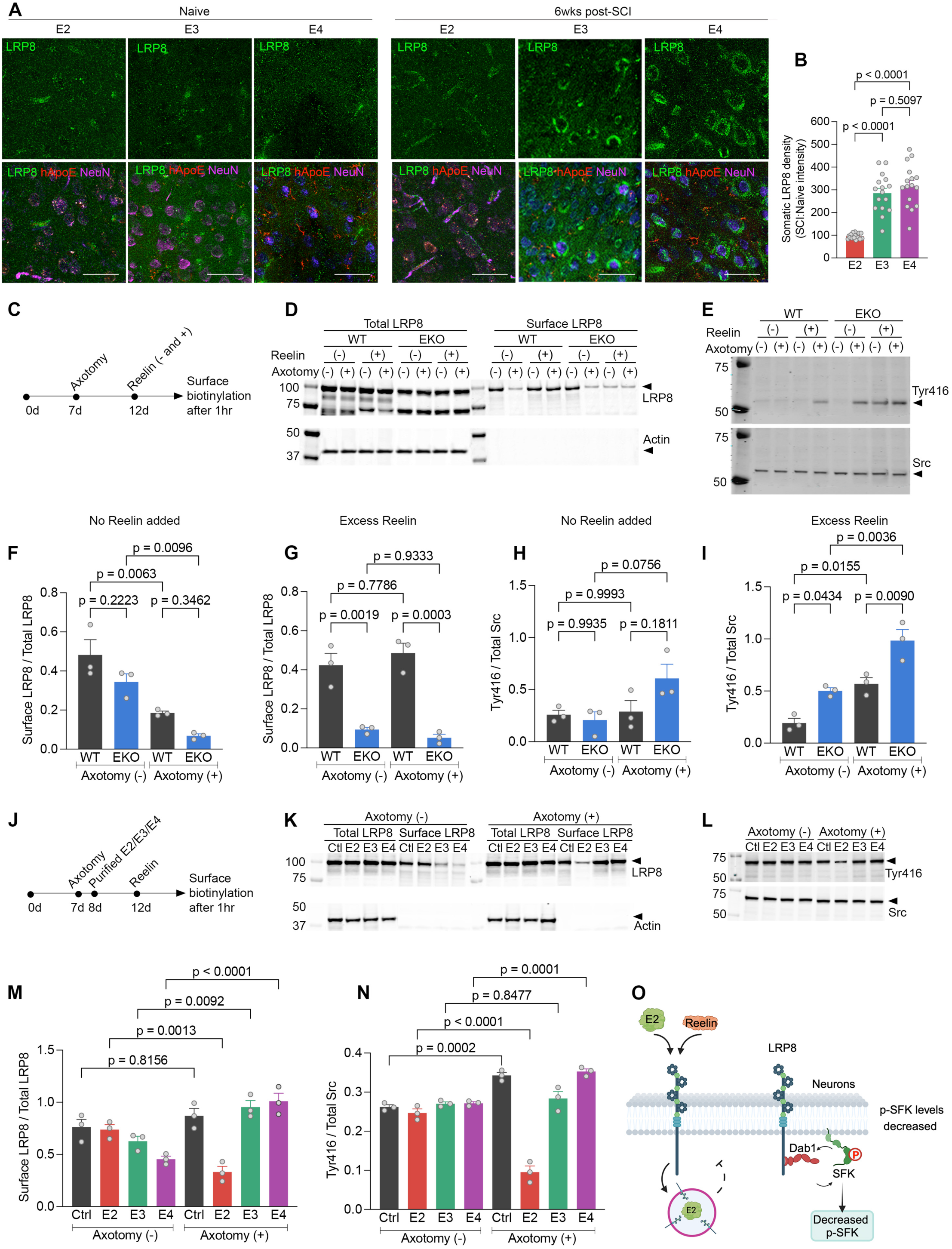
E2 impairs reelin-dependent recycling of LRP8 receptor and Src kinase activation in axotomized cortical neurons. (A) Photomicrographs of L5 from M1 brain cortex stained for anti-LRP8 (green), anti-hApoE (red) and anti-NeuN (magenta) 42 days after SCI compared to age-matched naïve controls for indicated genotypes. Scale bar, 200 µm. (B) LRP8 levels in neuronal soma at 42 days after SCI for E2, E3, E4 mice (*n* = 14). (C) Overview of study design to test influence of axotomy and reelin in LRP8 recycling and pTyr416-Src Family Kinase (SFK) activation in WT and EKO neurons. (D) LRP8 immunoblot using total and biotinylated surface protein extracts from (C) with actin as control. (E) pTyr416-SFK immunoblot using extracts from (C) with total Src as control. (F) Surface LRP8 with no added reelin from three independent experiments described in (C). (G) Surface LRP8 with excess reelin from three independent experiments described in (C). (H) Src kinase activation with no added reelin from three independent experiments described in (C). (I) Src kinase activation with excess reelin from three independent experiments described in (C). (J) Overview of study design to test influence of purified allelic ApoE lipoparticle preparations and reelin on LRP8 recycling and pTyr416-SFK activation in axotomized WT cortical neurons. (K) LRP8 immunoblot using total and biotinylated surface protein extracts from (J) with actin as control. (L) pTyr416-SFK immunoblot using extracts from (J) with total Src as control. (M) Surface LRP8 with excess reelin from three independent experiments described in (J). (N) Src activation with excess reelin from three independent experiments described in (J). (O) Schematic of cellular events in regenerating WT cortical neurons treated with E2. LRP8 undergoes rapid endocytosis in presence of reelin. Astrocytic E2 sequesters LRP8 to intracellular compartments thereby reducing downstream SFK activation in regenerating cortical neurons. Data shown as mean ± SEM and *p* values calculated by one-way ANOVA with Tukey’s multiple comparisons test for (B): F = 43.74, dF = 2; for (F): F = 16.36, dF = 3; for (G): F = 29.30, dF = 3; for (H): F = 3.47, dF = 3; for (I): F = 24.02, dF = 3; for (M): F = 15.82, dF = 7; for (N): F = 59.86, dF = 7.

We next evaluated LRP8-related signal transduction in cultured neurons as a function of axotomy and ApoE, focusing on the status of receptor-dependent Src family kinase activation as well as cell surface LRP8 localization. We performed a cell surface biotinylation experiment in wild-type and EKO primary cortical neurons to assess surface LRP8 retention while monitoring phospho-Src family (pTyr416) levels (Fig. 4C). Cytoplasmic Src kinase activity is regulated by phosphorylation of Y527 and Y416. Y527 suppresses kinase activity whereas Y416 enhances kinase activity by stabilizing the activation loop for substrate binding. Therefore, we monitored pTyr416 levels. In WT neurons, axotomy alone led to internalization of more than 50% of LRP8 (Fig. 4D and 4F), with little change in Src activation (Fig. 4E and 4H). Deletion of ApoE synergized with the axotomy-induced LRP8 internalization, and led to increased pTyr416 levels. Since both Reelin and ApoE are ligands regulating LRP8, we included experiments with exogenous Reelin added. In WT neurons, excess Reelin prevented axotomy-induced LRP8 internalization (Fig. 4D and 4G), but supported axotomy-induced Src activation (Fig. 4E and 4I). In ApoE null cultures with Reelin added, surface levels of LRP8 were low even without axotomy, and axotomy-induced Src activation was enhanced synergistically to the highest level. Thus, there is a robust interaction of axotomy, excess reelin and ApoE deletion with regard to LRP8 internalization and Src kinase activation.

We considered how ApoE isoforms might interact with axotomy to regulate LRP8 and Src in cortical neurons. It is known that ApoE isoforms differ in their binding to LRP receptors(*25, 30, 31*) and in their regulation of synaptic homeostasis in AD(*32–34*). Therefore, we examined the effect of different purified ApoE lipoparticles on the axotomy response for both Src activation and LRP8 internalization in wild-type cortical neurons supplemented with Reelin (Fig. 4J). Without injury, excess human ApoE did not substantially alter surface LRP8 level or Src activation (Fig. 4K, 4M and 4N). With no exogenous human ApoE particles added, axotomy elevated pTyr416 phospho-Src family levels moderately, but did not alter LRP8 distribution. In the presence of added E2, axotomy reduced surface LRP8 and strongly suppressed Src activation. In stark contrast, cultures with E4 showed increased surface LRP8 and Src activation after axotomy. The effects of E3 were intermediate. In contrast to LRP8, surface localization of LRP5 and LRP6 was not regulated by axotomy of ApoE (Fig. S4E-S4H). These data demonstrate strong interactions between axotomy and ApoE isoforms to regulate LRP8 surface levels specifically and to couple with downstream Src signaling.

### Forebrain excitatory neurons show an ApoE allele-specific transcriptomic response to spinal injury

We sought to provide a comprehensive transcriptomic view of forebrain responses to SCI as a function of ApoE genotype given the evidence for greater repair and recovery in E2 mice coupled with altered LRP8 signaling. We profiled single cell transcriptomes in ApoE knock-in mice with and without SCI at a relevant time point to uncover pro-regenerative signatures. Dorsal over hemi-sections of the thoracic spinal cord of E2 and E4 mice were created, as in the behavioral and tracing studies above. Single nuclei were extracted from the forebrain region of E2 and E4 mice without SCI or at 11 dpi (days post-injury) for snRNAseq. Data from 20 mice across the four groups (E2, E4, E2-SCI, E4-SCI) passed our quality check and were used for cellular profiling. Data filtering and cell clustering retained 201,941 nuclei for assessment of cellular composition and expression (Fig. S5). To create a cellular pro-regeneration atlas, our clustering paradigm segregated 10 major cell classes consistent with standard markers: ExNeurons, InNeurons, OPCs, oligodendrocytes, astrocytes, microglia, pericytes, vLMCs, endothelial, ependymal. Within clusters, the nuclei frequencies for the four experimental groups were similar (Fig. S5). This is consistent with limited cell death or proliferation in the forebrain distant from the SCI site at this timepoint. We therefore focused on gene expression changes within cell clusters that might correlate with the differential response to injury and ApoE allele.

To evaluate E2-specific pro-regenerative genes, we identified differentially expressed genes as SCI-induced (SIG; E2 versus E2-SCI) or regeneration-associated (RAG; E4-SI versus E2-SCI), and then focused on their intersection as SIG-RAG genes (Fig. 5). This yielded 320 SIG-RAG genes in ExNeurons and 248 in InNeurons. Strikingly, there were far fewer SIG-RAG genes in any glial cell type. While 49 and 68 genes in the SCI-RAG were identified for oligodendrocytes and astrocytes, respectively, only a single gene met these criteria in microglial cells. Thus, the primary molecular changes in the forebrain associated with greater recovery and repair in the E2-SCI mice are neuronal, and not glial. In excitatory neurons, a heatmap of SCI-RAG genes (Fig. 5C) showed that the transcriptomic response of the forebrain to SCI was largely opposite for E2 versus E4. The expression of many genes was induced in forebrain ExNeurons of the E2-SCI mice, while being suppressed in E4-SCI mice. The few genes induced in E4-SCI neurons, were suppressed in E2-SCI mice. These profiles provide a molecular correlate of the differential repair and recovery outcomes of these mice from SCI.

**Fig. 5.**
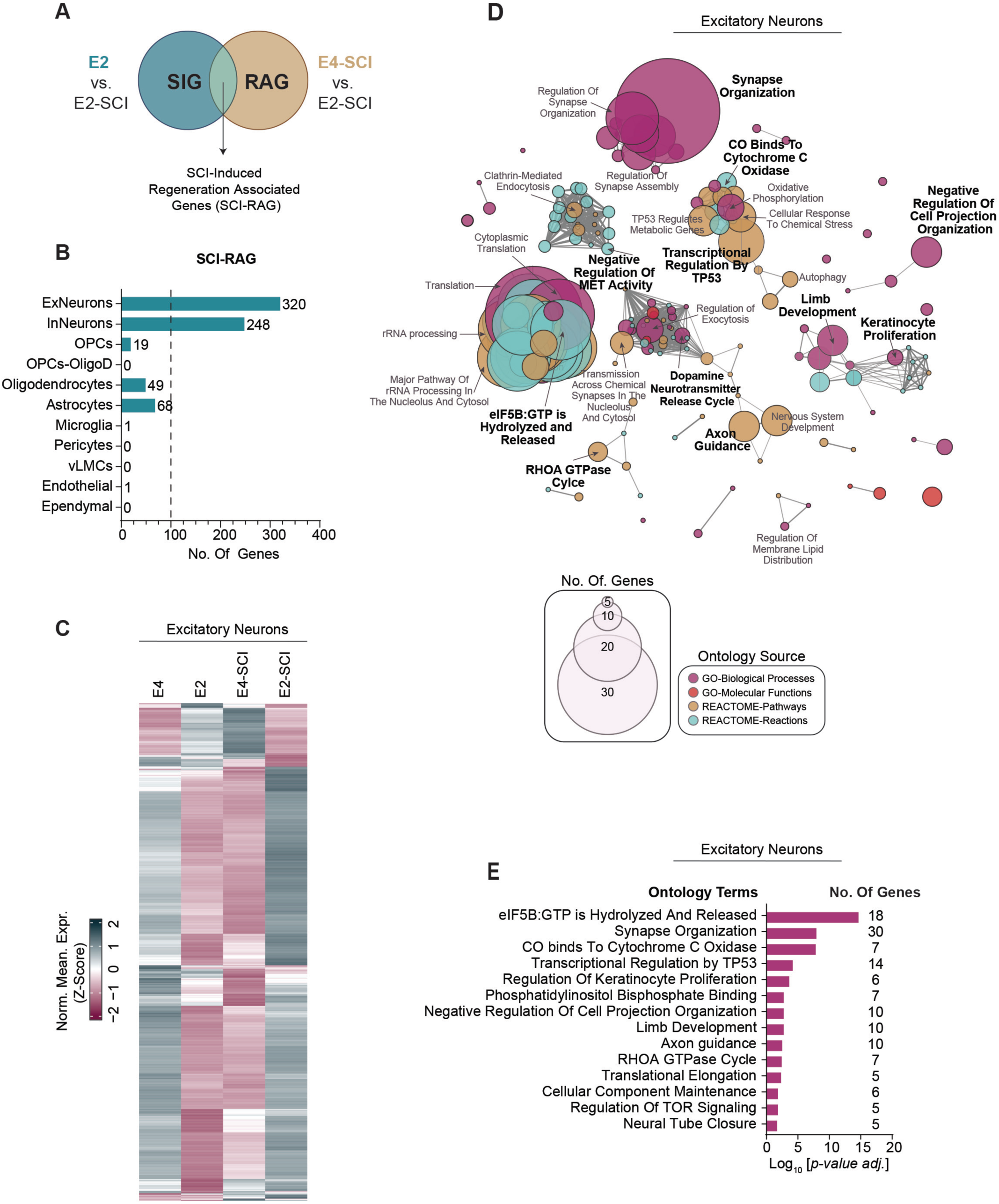
SCI-induced forebrain gene expression changes dependent on ApoE and linked to axon regeneration. (A) Schematic describing comparative analysis to identify SCI-induced regeneration-associated genes (SCI-RAGs. Genes are deemed as SCI-RAGs if they are differentially expressed post-SCI (E2 vs E2-SCI) and are associated with the post-SCI regenerative phenotypes of E2 (E4-SCI vs E2-SCI). (B) Gene count of identified SCI-RAGs across multiple cell types. The identified SCI-RAGs for pooled excitatory and inhibitory neurons listed in Supplemental Table S1. (C) Heatmap showing the relative normalized expression of SCI-RAGs of excitatory neurons. (D) Pathway enrichment analysis of excitatory neurons SCI-RAGs depicted in B and C. Enrichment terms (nodes) are organized into functional groups linked by shared gene associations (edges). Nodes are color-coded according to their respective gene ontology (GO) source and scaled in size by the number of gene-term associations. Intra-group significant terms (leading terms), in bold, are selected based on their respective *p*-values. A complete list of enriched pathway terms can be found in Supplemental Table S2. (E) Bar plot of the leading terms, their *p-*values and associated gene counts, for the pathway enrichment analysis of excitatory neuron SCI-RAGs shown in D.

To characterize the cellular processes underlying reparative responses, the differentially expressed SIG-RAG genes of ExNeurons were assessed for pathway enrichment (Fig. 5D, 5E). The most prominent pathways related to neural plasticity, axon guidance and RNA metabolism. Of particular note, terms regarding synaptic organization and post-synaptic density may relate to plasticity of supraspinal systems contributing to recovery after SCI, while those related axon guidance and RHOA GTPase signaling pertain to axonal growth and neural repair. Regulation of rRNA processing may reflect a general contribution of translational regulation to the E2-SCI phenotype.

Separate analysis of the SCI category and the RAG category yielded similar conclusions to the assessment of the SCI-RAG intersection (Suppl. Fig. S6). When comparing E2-SCI to E2 mice (Fig. S6A, S6B, S6D), the most prominent changes were expression increases in ExNeurons, with few glial changes. We observed that the E2-SCI mice showed some changes in endothelial gene expression relative to uninjured E2 mice that did not meet SCI-RAG criteria. In contrast, comparison of ExNeuron expression in E4-SCI vs E4 mice, showed predominantly down-regulated genes in the mice with poor SCI recovery (Fig. S6C). Pathway enrichment of ExNeuron E2-SCI vs E2 expression again identified synapse organization, GTPase regulation and translational regulation (Fig. S6D). We also assessed differential expression between E4 and E2 mice with SCI (Fig. S6E-G). As in the other comparisons, the most striking change was downregulation of genes in ExNeuron and InNeuron of E4 (upregulation in E2), with few glial changes. The major pathways identified amongst differentially expressed genes related to synapse organization plus neuronal development and projection. Thus, regardless of pairwise comparison strategy, the E2-SCI mice with enhanced repair and recovery exhibit expression of synaptic organization and axon projection genes in neurons that E4 mice fail to upregulate after SCI.

From the single cell transcriptomic profiles, we considered individual gene expression changes. The most robust individual gene changes of ExNeuron SCI-RAG genes identified amongst the pathway terms translation and synaptic organization are summarized in a heatmap (Fig. 6A). Notably, many of the synapse organization genes have evidence for protein-protein interactions with one another (Fig. 6B). We sought to validate a subset of these changes at the protein level in cerebral cortex by immunohistology and immunoblot using samples collected 6 weeks after SCI. The protein subset to be tested was based on possession of a known function and antibody availability. Immunohistological analysis of expression in NeuN-positive neurons of L5 in M1 cortex for Efnb2 and Synaptopodin matched the forebrain ExNeuron mRNA expression pattern with highest levels in E2-SCI mice (Fig. 6C-6E). This supports their potential role in modulating repair and plasticity in CST neurons of E2-SCI mice. The patterns for Tsc1 (Fig. 6C and 6F) and for Homer1 (Fig. 6C and 6G) showed statistically significant differences between groups with lower levels in E2-SCI than E4-SCI L5-M1 neurons. The differences between immunohistology and transcriptomics for Tsc1 and Homer1 may reflect changes unique to timing (6 weeks vs 11 days) or to region (L5-M1 vs anterior forebrain). Of note, the suppression of Tsc1 has been linked to the PTEN pathway and to increased axon regeneration in simple model systems(*13, 35*). Immunoblot of cortical protein levels were also completed for Tsc1, Efnb2, Homer1 and Synaptopodin (Fig. 6H-6M). The biochemical studies confirmed the regulation of these proteins by ApoE allele and SCI and confirmed the histological findings in all cases, with the exception of Homer1 in E4-SCI samples. Overall, the protein analysis verified the ApoE2- and SCI-dependent expression of axon and synapse regulating pathways in the cortical neurons relevant to SCI repair and recovery.

**Fig. 6.**
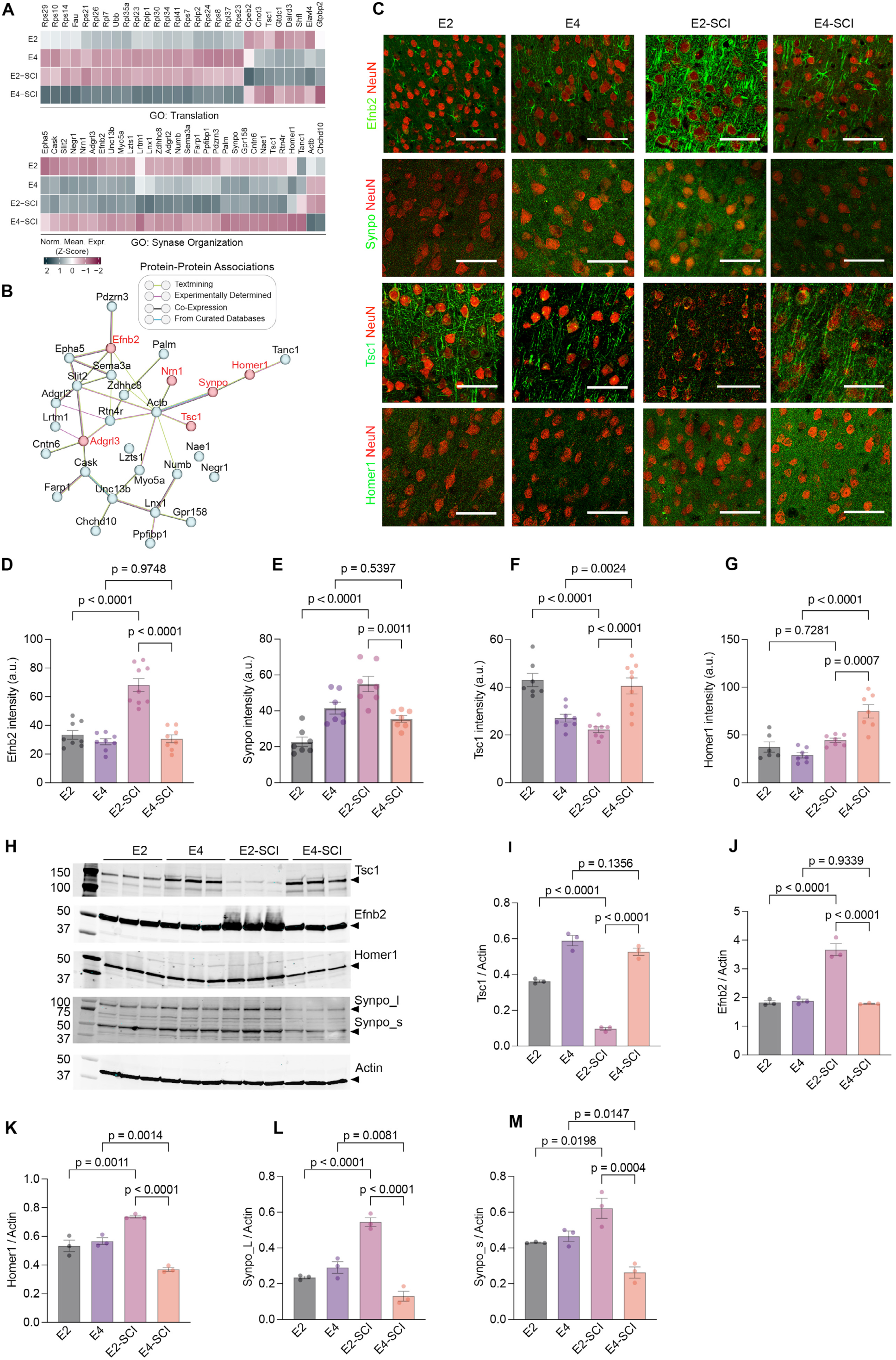
ApoE allele-specific regulation of synaptic protein expression in forebrain after SCI. (A) Heatmap showing normalized gene expression changes associated with translation and synapse organization GO terms from pathway enrichment analysis performed on excitatory neurons SCI-RAGs depicted in Fig. 5D and 5E, and listed Supplemental Table S2. (B) Protein interaction network of the GO: synapse organization associated genes shown in A. Protein associations (edges) are color-coded based to the evidence source of their protein-protein association. (C) Photomicrographs of L5 from M1 brain cortex stained for Efnb2, Synpo, Tsc1, and Homer1 in green and NeuN in red, 42 days after SCI compared to age-matched naïve controls for respective genotypes. Scale bar, 100 µm. (D) Mean signal intensity of Efnb2 in L5 of M1 cortex for E2 naive (*n* = 8), E4 naive (*n* = 8), E2-SCI (*n* = 9) and E4-SCI (*n* = 9) mice. (E) Mean signal intensity of Synpo in L5 of M1 cortex for E2 naive (*n* = 7), E4 naive (*n* = 7), E2-SCI (*n* = 7) and E4-SCI (*n* = 7) mice. (F) Mean signal intensity of Tsc1 in L5 of M1 cortex for E2 naive (*n* = 7), E4 naive (*n* = 8), E2-SCI (*n* = 8) and E4-SCI (*n* = 9) mice. (G) Mean signal intensity of Homer1 in L5 of M1 cortex for E2 naive (*n* = 6), E4 naive (*n* = 7), E2-SCI (*n* = 7) and E4-SCI (*n* = 7) mice. (H) Immunoblot analysis of selected top four candidates with total protein extracted from motor cortex and subcortical regions of respective genotype with actin as loading control. (I) Densitometry analysis of Tsc1 immunoblot. (J) Densitometry analysis of Efnb2 immunoblot. (K) Densitometry analysis of Homer1 immunoblot. (L) Densitometry analysis of Synpo_L immunoblot. (M) Densitometry analysis of Synpo_s immunoblot. Data shown as mean ± SEM. *p* values calculated by one-way ANOVA with Tukey’s multiple comparisons test for (D): F = 32.61, dF = 3; (E): F = 17.94, dF = 3; (F): F = 16.54, dF = 3; (G): F = 18.46, dF = 3; (I): F = 63.45, dF = 3; (J): F = 48.61, dF = 3; (K): F = 17.31, dF = 3; (L): F = 137.8, dF = 3; (M): F = 37.71, dF = 3.

### Therapeutic expression of E2 promotes neural repair in mice and human neurons

Given the prominent effect of ApoE alleles on repair, recovery and gene expression after SCI, we hypothesized that ApoE2 expression after SCI would confer a therapeutic benefit. We introduced AAV-ApoE2 versus AAV-GFP into wild type mice at 3 days after SCI using bilateral intraparenchymal injection into cerebral cortex to evaluate this intervention (Fig. 7A). Prior to SCI efficacy studies, we evaluated the level and distribution of ApoE2 expression by immunoblot and immunohistology in uninjured wild-type, EKO and E2 mice at 28 days post-injection. As detected by a human-specific ApoE antibody, the forebrain level of ApoE2 in AAV-ApoE2 injected wild-type, EKO was equivalent to the uninjected ApoE2 knock-in mice (Fig. S7A). For ApoE2 mice, the AAV injection increased forebrain protein level by about two-fold. We also assessed the distribution of human ApoE2 protein throughout the neuroaxis histologically. No staining signal was observed in uninjected wild-type or EKO samples, validating the specificity of the method. After cortical AAV-ApoE2 injection, there was prominent human ApoE in the cerebral cortex, similar to GFP expression from AAV-GFP (Fig. 7B). In addition, the secreted human ApoE2 protein diffused widely in the CNS with strong staining observed in spinal cord neurons where cortical AAV-GFP produced no GFP signal (Fig. S7B). Thus, cortical AAV-ApoE2 provided effective delivery of the protein broadly within the mouse CNS.

**Fig. 7.**
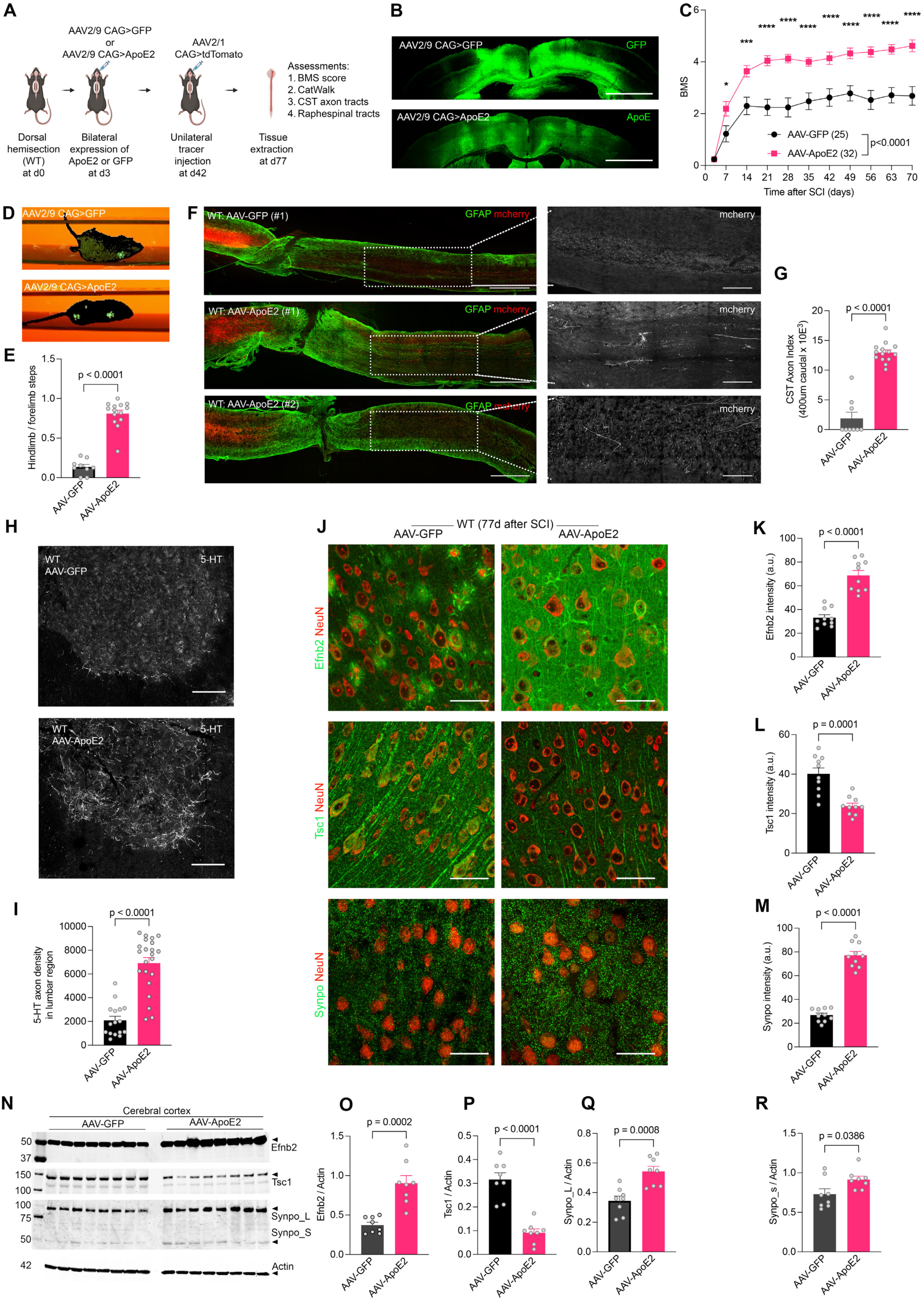
AAV-mediated expression of ApoE2 promotes functional and anatomical recovery of WT mice after SCI. (A) Overview of experimental approach to evaluate therapeutic benefit of ApoE2 in WT mice with dorsal thoracic over-hemisection injury. (B) Photomicrograph of brain cortex region of mice that received bilateral AAV2/9 injection into M1 cortex. Sections stained for anti-GFP and anti-hApoE at d77 post-injection. Scale bar, 500 µm. (C) BMS open-field locomotion assessment of WT-SCI mice receiving AAV-GFP (*n* = 25) and AAV-ApoE2 (*n* = 32). Performance scores were recorded for each animal every consecutive week. Data shown as mean ± SEM and analyzed using repeated measure multi-point ANOVA across time series followed by post-hoc Tukey’s multiple comparisons test between genotypes at indicated time points. ****p<0.0001, ***p<0.001, **p<0.01 significant difference between genotypes. (D) Single frame CatWalk video showing fore and hindlimb paw footprints in green for WT-SCI mice injected with AAV-GFP or AAV-ApoE2, and observed at d77 after SCI. (E) Limb coordination index for WT-SCI mice treated with AAV-GFP (*n* = 9) or AAV-GFP (*n* = 14), d77 after SCI. (F) Sagittal low-power photomicrographs of spinal cord around the lesion site in AAV-injected WT mice at d77 after SCI. Sections were stained with anti-GFAP (green) and anti-mCherry (red). Dorsal is up and rostral is left. Scale bar, 500 µm. White outlined boxed areas in each image are captured at high-resolution to visualize regenerating CST fibers caudal to lesion for red channel only. Scale bar, 100 μm. (G) Quantification of CST axon index in the lumbar spinal cord caudal to lesion. CST fiber numbers were measured at 400 µm caudal to the lesion site in WT mice expressing GFP (*n* = 9) or ApoE2 (*n* = 14) at 77 days after SCI. (H) Transverse axis photomicrograph of ventral horn lumbar spinal cord caudal to lesion stained with anti 5-HT. Scale bar, 100 µm. (I) Quantification of 5-HT+ve serotonergic fiber density in ventral horn of lumbar spinal cord for AAV-GFP (*n* = 16) and AAV-E2 (*n* = 22) mice at d77 after SCI. (J) Photomicrographs of L5 from M1 brain cortex stained for Efnb2, Synpo, Tsc1 in green and NeuN in red, for WT-SCI mice treated with AAV-GFP (*n* = 10) and AAV-ApoE2 (*n* = 10). Scale bar, 100 µm. (K) Mean signal intensity of Efnb2 in L5 of M1 cortex for therapeutic WT-SCI mice. (L) Mean signal intensity of Tsc1 in L5 of M1 cortex for therapeutic WT-SCI mice. (M) Mean signal intensity of Synpo in L5 of M1 cortex for therapeutic WT-SCI mice. (N) Immunoblot of selected proteins in extracts from cerebral cortex of respective genotypes with actin as loading control. (O) Densitometry analysis of Efnb2 immunoblot. (P) Densitometry analysis of Tsc1 immunoblot. (Q) Densitometry analysis of Synpo_L immunoblot. (R) Densitometry analysis of Synpo_s immunoblot. Data shown as mean ± SEM. *p* values calculated by two-tailed unpaired t-test for (E): t = 10.9, df = 21; (G): t = 9.7, dF = 11; (I): t=3.22, df=13; (K): t = 7.3, df = 18; (L): t = 4.8, df = 18; (M): t = 13.6, df = 18; (O): t = 5.4, df = 14; (P): t = 6.9, df = 14; (Q): t = 8.8, df = 14; (R): t = 4.2, df = 14.

The functional role of human ApoE2 overexpression was assessed in adult wild-type mice with thoracic dorsal over-hemisection injury, the same traumatic lesion utilized for the ApoE knock-in and knockout studies described above. On the third day after SCI, all mice were incapable of locomotor function with BMS scores of 1 or less. The animals were randomized to receive AAV-ApoE2 or control AAV-GFP virus at that point. The degree of locomotor recovery as measured by BMS in the open field was significantly greater in the AAV-ApoE2 group (at 70 days, 4.70 ± 0.24, mean ± SEM, n=32) than in the AAV-GFP group (2.44 ± 0.36, n=25) (Fig. 7C). Nearly all AAV-ApoE2 showed occasional or consistent plantar stepping with weight support, while most control AAV-GFP mice exhibited ankle movement but not plantar stepping or weight support. In CatWalk analysis of walking along a linear track, both limb coordination and hindpaw footprints were consistently detected in ApoE2-treated compared to GFP group (Fig. 7D). Automated analyses detected hindlimb steps for over 75% of forelimb cycles in the AAV-ApoE2 group, while recognizing hindlimb steps associated with fewer than 10% of forelimb steps by the AAV-GFP mice (Fig. 7E). After 8 weeks survival, spinal cords were collected for histology. The E2-treated group showed a greater than four-fold increase in caudal CST fibers (Fig. 7F-7G) and two-fold increase of raphespinal axon length in the ventral horn of lumbar spinal cord compared to GFP-treated group (Fig. 7H and 7I). Multiple additional examples are shown in Fig. S7C-S7D).

We sought to confirm whether AAV-ApoE2 expression after SCI generated the same molecular profile as observed for constitutive ApoE2 knock-in SCI mice. The differentially expressed proteins from SIG-RAG analysis of knock-in mice were assessed in the therapeutic AAV cohort by immunohistology and immunoblot. Localization (Fig. 7J-7M) and cortical expression (Fig. 7N-7R) changes of Efnb2, Tsc1 and Synpo in NeuN-positive L5 neurons of M1 cortex matched with the ApoE2 knock-in expression changes. These results highlight the importance of axon and synaptic plasticity pathways for therapeutic benefit of ApoE2 after SCI. Overall these results demonstrate post-injury delivery of an ApoE2 based therapeutic improves SCI recovery and neural repair.

We considered whether human neurons as well as mouse neurons were responsive to the pro-regenerative benefit of ApoE2. We examined axon regeneration from iPSC-derived glutamatergic neurons derived from stable integration of neurogenin-2 under doxycycline-inducible promoter (i^3^Neurons)(*36*). Twenty-four hours after axotomy of mature neurons on the 50^th^ day post differentiation, AAV-E2 or AAV-GFP was added to these *APOE3* homozygous neuronal cultures (Fig. S7G). A substantial increase of both cellular and secreted ApoE protein was confirmed for the neurons treated with AAV-E2 (Fig. S7E and S7F). Axon regeneration over the subsequent 7 days was twice as extensive for the cultures expressing ApoE2 as compared to those expressing GFP (Fig. S7H-S7I). These data support the potential therapeutic benefit for ApoE2 overexpression after CNS trauma.

## DISCUSSION

The central finding of this study is that neurons exposed to secreted ApoE2 exhibit regenerative axon growth after axotomy, a process dependent on LRP8-mediated signaling. In contrast, exposure to ApoE4, or lack of ApoE, limits axon regeneration. After spinal cord injury, the presence of ApoE2 before or after trauma supports greater corticospinal and raphespinal axon growth, improved behavioral recovery, and altered neuronal transcriptomes. Axotomy and reelin trigger LRP8 redistribution and downstream kinase activation, while ApoE2 antagonizes these effects and supports injury-induced upregulation of synapse organization genes.

A broad range of pathways have been implicated in axon regeneration using various model systems(*37–42*). ApoE transcription is known to modulate dorsal root ganglion sensory axon outgrowth(*43*). In contrast, genetic linkage studies in humans tracking recovery after traumatic CNS injury have thus far identified only a few genetic variations influencing repair or recovery(*19, 20, 44, 45*). Here, we investigated the ApoE–LRP8 axis, since the LRP8 receptor was a candidate from our *in vitro* axon regeneration screening(*10*), and genetic variation in its ApoE ligand has been linked to the outcome of human TBI recovery(*19*). Convergent discovery of this ligand receptor pair across the translational spectrum emphasizes the potential importance of these molecules.

The ApoE action studied here appears to be selective for neurons. The culture system includes predominantly neurons and secreted ApoE from astrocyte cultures transferred activity to ApoE knockout neuronal cultures. *In vivo* spinal cord injury studies showed robust effects on regenerative axon growth and pronounced interactions of spinal cord injury and ApoE alleles on neurons distant from the trauma site. It is abundantly clear from studies of neurodegeneration that ApoE alleles differentially affect glial cells, with alterations in microglial and astroglial metabolism, reactivity and function(*29, 32, 33, 46–49*). In addition, ApoE regulates macromolecular transport and metabolism to participate in neurodegenerative mechanisms(*50*). In the studies here, while the source of ApoE is likely to be predominantly glial, there is no evidence that glial reactions at the injury site or in the forebrain provide a substantial contribution to the improved axon growth and functional recovery. In particular, tissue sparing and glial reaction were not altered by different ApoE alleles. Many fewer differentially expressed genes were observed for microglia and astrocytes than for neurons in the brain after injury. Although ApoE’s most studied effect is on Aß accumulation and Alzheimer’s risk, our SCI and axon regeneration models show no evidence for Aß accumulation or altered APP.

The analysis of LRP8 subcellular localization and signal transduction delineate ApoE allele specific actions. Reelin and axotomy share actions to decrease surface LRP8 and increase Src family kinase activation. ApoE2, but not ApoE4 or ApoE3, antagonizes the action of axotomy and excess reelin. These data are consistent with previous descriptions of antagonistic action of ApoE and reelin for Lrp8 without axotomy(*51*). SCI in the presence ApoE2 leads to increased expression of multiple genes and proteins associated with synaptic organization. Gene products such as Efnb2, Synaptopodin, and Homer1 are likely to contribute to neural repair. Their increased expression is consistent with the formation of new and plastic synaptic connections. The suppression of Tsc1 in ApoE2 – SCI mice is consistent with an increased intrinsic axon regenerative growth state(*13, 35*). Overall, our SCI and transcriptomic studies highlights a broad cellular heterogeneity of forebrain neurons in synaptic modulation of spinal circuits for recovery in response to secreted ApoE along the neuroaxis. The relative importance of different circuits is not studied here, but likely many different neuronal populations participate in addition to corticospinal and raphespinal neurons.

One implication of these studies is that ApoE genotype can alter the prognosis for recovery from CNS trauma, likely both SCI and TBI. Worse outcomes for ApoE4 cases of SCI have been reported(*20–23*), though a potentially beneficial effect for the rarer ApoE2 allele has not been described. Consideration of ApoE genotyping to increase prognostic accuracy and to stratify clinical trials of various interventions may be warranted(*52*).

We utilized AAV injection to overexpress ApoE2 in wild type mice beginning three days after injury in a therapeutically relevant design and observed improved function and axon growth. Future experiments with contusive injury, longer delays to treatment, and systemic vectors will explore the interventional potential of this approach. In addition, kinases downstream of Lrp8 may be critical for ApoE2 benefit such that small molecule inhibitors may provide alternate sites for intervention.

These findings also have important implications for the role of ApoE in neurodegeneration. ApoE expression, and ApoE2 in particular, may be expected to promote compensatory mechanisms, through neuronal plasticity and axonal sprouting, to mitigate symptoms in Alzheimer’s disease and related disorders. Specifically, ApoE genotype may modulate progression rate in Alzheimer’s disease via compensatory mechanisms unrelated to the risk of disease or the accumulation of Aß and tau or inflammation. Thus, the role of ApoE genotype in neuronal resilience may be a critical, but still understudied, aspect of neurodegeneration. In this regard, our data from axotomy studies indicate that ApoE2 will be most beneficial while ApoE4 is most detrimental. The spinal trauma model indicates that suppression of all ApoE expression is deleterious for compensatory repair after CNS damage.

It is known that many cases of dementia are multifactorial, with vascular disease frequently co-occurring with Alzheimer’s pathology. Disruption of neuronal connectivity due to vascular compromise of white matter may respond to ApoE genotype in the same manner that disconnection due to spinal cord trauma does. Regardless of whether neuronal connectivity is damaged by vascular insufficiency, neurodegenerative protein accumulation or trauma, we expect ApoE signaling to be a critical determinant of the success of neural repair and symptomatology.

In conclusion, ApoE2 has a robust effect to increase adult CNS axon growth after trauma that is dependent on altered LRP8 signaling with increased expression of synapse organizing genes. ApoE action is predominantly neuronal in this condition. Allele-specific ApoE expression alters functional recovery after traumatic SCI in mice, with overexpression of ApoE2 improving locomotor recovery.

## AUTHOR CONTRIBUTIONS

Conceptualization, R.K. and S.M.S.; Methodology, R.K., X.W., N.L., E.M.H., A.B., Y.S. and S.M.S.; Investigation, R.K., X.W., N.L., E.M.H., A.B., and I.I.; Writing – Original Draft, R.K. and S.M.S.; Writing – Review & Editing, all; Funding Acquisition, S.M.S.; Resources, S.M.S; Supervision, S.M.S.

## ACKNOWLEDGMENTS

This work was supported by grant R35NS097283 from the N.I.H. to S.M.S.

## SUPPEMENTARY MATERIALS

### METHODS

#### Animals

Spinal cord injury study was performed in mice with following genotypes - C57BL/6J (JAX stock#000664), ApoE KO (JAX stock #002052), ApoE2 KI (JAX stock #029017), ApoE3 KI (JAX stock #029018), ApoE4 KI (JAX stock #027894) and Lrp8 KO(*53*) (JAX stock #003524). Mice were maintained on a 12-h light/dark cycle with regular food and water under 40%-60% humidity. All animal procedures were performed following institutional guidelines and regulations. To minimize investigator bias, the study was blinded during entire analysis.

#### Primary neuronal cultures

Primary mouse cortical neurons were cultured in Neurobasal supplemented with B27 (10X) and L-glutamine (2 mM) while iPSCs were maintained in Essential 8 Medium (Gibco A1517001) and iPSC-derived glutaminergic human neurons were maintained in DMEM/F-12 based medium supplemented with essential growth factors described in detail below. All cells were maintained at 37°C with 5% CO_2_.

#### Cell culture and transfection

HEK293T (ATCC, CRL11268) cells were cultured in DMEM/F-12 (Gibco 11320-033). To ensure absence of mycoplasma, all cell lines were periodically examined using LookOut Mycoplasma PCR kit (Sigma-Aldrich, MP0035). A day before transfection, HEK293T cells were passaged at a density of 10^7^ cells/15-cm plate and transfected at 80% confluency with respective constructs for AAV production. DNA: PEI transfection mixture containing 150 µl of polyethylenimine (PEI, Polysciences Inc.) incubated at RT for 15 min before adding to HEK293T cells. All cells were maintained at 37°C with 5% CO_2_.

#### Mouse behavioral tests

Two researchers unaware of the mice genotype performed all behavioral tests. We used Basso Mouse Scale (BMS) as a measure of open-field locomotion(*54*). BMS has a quantitative scale from 0 to 9. BMS scoring were done once pre-injury and starting on 3d of injury and weekly thereafter for all dorsal thoracic over-hemisection experiments.

The CatWalk XT (v 10.6) gait analysis system (Noldus, Netherlands) design consisted of a 1.5 m black corridor walkway on a black glass plate using green LED bottom lighting. Entire system is placed in dark silent room. Paw prints were captured by 100 fps high-speed camera positioned beneath the black glass floor. Walkway area and compliance parameters are set following company recommendations. Mice were habituated to the system and the walkway voluntarily, twice a day for three days to complete three accomplished runs. At the end of training period, mice were tested by crossing the corridor three times. To detect the paws from the background, we used same detection settings for all genotypes (camera gain: 20.50, green walkway light (15.5, green intensity threshold: 0.15, red ceiling light :17.3). To measure the limb coordination, we measured fore-hind step ratio using number of fore and hind limb steps taken by each mouse across three replicate runs.

#### Surgical procedures for SCI, CST tracing and CNS gene therapy experiments

All animal procedures and post-operative care were performed in accordance with the Institutional Animal Use and Care Committee guidelines at Yale University. Age-matched adult (12-13 weeks) female mice were subjected to dorsal thoracic over-hemisection (75% depth) as described previously(*55*). All animals received subcutaneous injection of 100 mg/kg ampicillin and 0.1 mg/kg Buprenex twice a day for the first 2 d after surgery and additional injections later as necessary.

To trace CST tracts, we unilaterally injected AAV9-CAG>tdTomato (Addgene #59462) into sensorimotor cortex to anterogradely label the CST at each of the five sites (coordinates from bregma in mediolateral/anterior–posterior format in mm: 1.0/0.0, 1.5/1.5, 1.5/0.5, 1.5/−0.5, 1.5/−1.5) for a total of 1.5 µl volume. Mice were kept for additional 4 weeks before being euthanized for morphometric analysis. For expression of ApoE2 gene therapy experiments, AAV9-CAG-ApoE2 and control AAV9-CAG-GFP (Addgene #37825) was injected bilaterally into sensorimotor cortex (coordinates as mentioned above) at 3 d after dorsal hemisection procedure. All animals underwent surgery received subcutaneous injection of 100 mg/kg ampicillin and 0.1 mg/kg Buprenex twice a day for the first 2 d after surgery and additional injections later as necessary. The study was randomized and blinded for genotypes and virus used.

#### AAV-ApoE2 vector construction and AAV9-ApoE2 virus production

The coding sequence for human ApoE2 was amplified from pCMV4-ApoE2 (Addgene#87085) and inserted at BamH1 and HindIII sites of pAAV-CAG (Addgene#59462) upstream of WPRE and SV40 pA sequence by replacing tdTomato coding region in #59462. ApoE2 fragment was amplified using Q5^R^ Hot Start High-Fidelity 2X master mix (# M0491L) according to manufacturer protocol with ApoE-specific primers: forward primer (20mer): ATGAAGGTTCTGTGGGCTGC and reverse primer (20mer): GCAGCCCACAGAACCTTCAT. The final vector pAAV-CAG>hApoE2-WPRE-SV40pA was sequence verified in-house using internal and vector specific sequencing primers.

A day before transfection, HEK293T cells were passaged at a density of 10^7^ cells/15-cm plate and transfected at 80% confluency. For AAV production, the plasmid cocktail containing - pDF6 helper (18 µg of Addgene#112867), pAAV-CAG>ApoE2 (6 µg), AAV packaging plasmid expressing Rep/Cap genes pAAV2/9n (6 µg of Addgene#112865) were prepared in 3 ml of serum-free DMEM. DNA:PEI transfection mixture containing 150 µl of polyethylenimine (PEI, Polysciences Inc.) was incubated at RT for 15 min before adding to HEK293T cells. After 96 h following transfection, the cells were harvested and treated with DNase1 (10 U/ml, AMPD1, Sigma-Aldrich) and benzonase (50 U/ml, #70746 Millipore) for 40 min at 37°C. The mixture was centrifuged at 3,000 x *g* for 20 min at 4°C to remove cell debris. The supernatant containing viral particles were further purified by ultra-centrifugation using iodixanol density gradient(*56*). To estimate the viral titers samples were compared against a standard curve generated from known titer diluted from 10^8^ to 10^13^ genome copies per ml. Viral titers were determined using iQ SYBR Green Supermix (Bio-Rad) and quantitative PCR (Bio-Rad CFX96). Average AAV9-ApoE2 viral titers used for *in vitro* transduction of iPSC derived human neurons were 10^10^-10^11^ genome copies per ml and 10^11^-10^12^ genome copies per ml for *in vivo* CNS transduction experiments.

#### Primary mouse cortical neuron culture

We used postnatal d1 pups (P1) from respective genotypes to dissect cortices in ice-cold BrainBits medium. Enzymatic dissociation in 1X HBSS (Mg/Ca-free) supplemented with 100 U/ml DNAse (04716728001, Roche) containing Papain (20 U/ml, Worthington LK003178), 0.5M EDTA and 1 mM CaCl_2_ at 37°C for 30 min followed by mechanical dissociation. Cells were counted using hemocytometer and diluted to achieve a seeding density of 2.5-4.0 x 10^4^ per 200 µl for each well and plated on Corning BioCoat poly-D-Lysine coated glass 96-well plates (#354461) for axon regeneration assays and 12-well plates (#354470) for biochemistry experiments. Cultures were kept in 37°C with 5% CO2 incubator in Neurobasal-A media supplemented with 0.5% penicillin/streptomycin, 0.5% B27 and 2 mM Glutamax.

#### iPSC maintenance and differentiation of glutamatergic human neurons

Engineered iPSC expressing mammalian NGN2 (neurogenin 2) under doxycycline-induced system in the AAVS1 safe harbor locus, termed as GMK2 iPSC cell line also express inducible CRISPR interference machineries (pC13N-dCas9-BFP-KRAB). The parental human iPSC cell line used was CRISPRi-i3N iPSCs(*36*). The aliquots of the frozen cells in clumps were thawed at 37°C water bath and resuspended in Essential 8 Medium (GIBCO/Thermo Fisher Scientific; Cat. No. A1517001) containing 10 nM ROCK inhibitor Thiazovivin (#72254, Stem Cell Technologies) [E8+T]. iPSCs were centrifuged 350 x *g* at 4°C. The pelleted cells were resuspended in [E8+T] and platted in 1-well of a Vitronectin (#A31804, Vitronectin (VTN-N) Recombinant Human Protein, Truncated, GIBCO /ThermoFisher) coated 6-well plate. Cells were allowed to grow for 3-4 d or at 70-80% confluency to form the iPSC clumps with everyday [E8+T] media exchange. 70-80% iPSC cell culture was then split with Gentle Dissociation reagent (GDR) and again plated as clumps containing E8 medium in the vitronectin coated 6 well plates. In this way the iPSCs were maintained in a pluripotent state.

iPSCs were then allowed to grow (70-90%) confluency to differentiate and induce iPSCs into glutamatergic neurons. iPSCs were released by incubating the cells at *37*°C for 7 mins with StemPro Accutase Cell dissociation reagent (GIBCO Cat No: A11105-01) and centrifuged at 250 x *g* at 4°C for 5 mins. Next, the pelleted iPSC cells were resuspended in N2 pre-differentiation medium containing knockout DMEM/F12 (GIBCO/Thermo Fisher Scientific; Cat. No. 12660-012), 1X MEM non-essential amino acids (GIBCO/Thermo Fisher Scientific; Cat. No. 11140-050), 1X N2 supplement (GIBCO/Thermo Fisher Scientific; Cat. No. 17502-048), 10 ng/mL NT-3 (PeproTech; Cat. No. 450-03), 10 ng/mL BDNF (PeproTech; Cat. No. 450-02), 1 μg/mL mouse laminin (Thermo Fisher Scientific; Cat. No. 23017-015) and 10 nM ROCK inhibitor Thiazovivin (#72254, Stem Cell Technologies) to induce mNGN2 expression. Pelleted iPSCs were counted and plated at 7 x 10^5^ cells per well in a matrigel-coated 6-well plate contained 2 ml of N2 pre-differentiation medium per well. iPSC cells were grown in single cells for three days in N2 pre-differentiation medium. Three days later, as d0, the pre-differentiated cells were released by incubating the cells at 37°C for 7 mins with StemPro Accutase Cell Dissociation Reagent (GIBCO Cat No: A11105-01) and then centrifuged at 250 x *g* at 4°C for 5 mins. The pelleted pre-differentiated cells were then resuspended in classic neuronal medium containing the following: half DMEM/F12 (GIBCO/Thermo Fisher Scientific; Cat. No. 11320-033) and half Neurobasal-A (GIBCO/Thermo Fisher Scientific; Cat. No. 10888-022) as the base, 1X MEM Non-Essential Amino Acids, 0.5X GlutaMAX Supplement (GIBCO/Thermo Fisher Scientific; Cat. No. 35050-061), 0.5X N2 Supplement, 0.5X B27 Supplement (GIBCO/Thermo Fisher Scientific; Cat. No. 17504-044), 10 ng/mL NT-3, 10 ng/mL BDNF, 1 μg/mL Mouse Laminin, and 2 μg/mL doxycycline hydrochloride. Pre-differentiated cells were counted and plated at (5,000-10,000) cells per well of a BioCoat Poly-D-Lysine 96-well plate (Corning; Cat. No. 356640) in 100 µl of Classic Neuronal Medium per well. After 7 DIV, half of the classic neuronal medium was removed, and an equal volume of fresh medium was added without doxycycline. At 14 DIV, half of the medium was removed and twice that volume of fresh medium without doxycycline was added. At 21 DIV, one-third of the medium was removed and twice that volume of fresh medium without doxycycline was added. At 28 DIV and every week after, one-third of the medium was removed and an equal volume of fresh medium without doxycycline was added. In this way the neurons were allowed to grow for 45-50 days.

#### Astrocyte conditioned medium (ACM)

Wild-type, EKO and E2/E2/E4 KI mouse cerebral cortices were removed from postnatal d1 pups (P1) in ice-cold BrainBits medium. McCarthy and deVellis (MD) astrocytes protocol was adopted for culturing MD astrocytes(*57*). Enzymatic digestion performed with 2 ml of 0.25% trypsin at 37°C for 20 min followed by 1 ml of DMEM+FBS to deactivate trypsin. After adding 0.05 mg/ml DNase, the tube was spun at 300 x *g* for 5 min and the cell pellet was re-suspended in 1 ml of fresh DMEM+FBS. This suspension was then filtered through 70 µm cell strainer and the cells were plated onto an uncoated plastic T-25 flask maintained in a rotary shaker at 200 rpm. After 3 DIV, old medium was replaced with astrocytes conditioned medium (ACM) containing 50% neurobasal + 50% DMEM without phenol red, glutamine, pyruvate, N-acetylcysteine (NAC) and penicillin-streptomycin. Cultures were then maintained for additional 5 d to enrich secreted factors from astrocytes. Thereafter, any dead cells and debris were removed by centrifugation at 3,000 x *g* for 30 min. ACM was concentrated with a 3kDa MWCO (Amicon, 15ml) at 3,000 x *g* for 15 min at 4°C. After Bradford estimation of total protein concentration in ACM, aliquots were stored at -80°C for further experiments.

#### Isolation of native ApoE lipoprotein particles from ACM

We coupled a mouse monoclonal WUE-4 apoE antibody that detects the three isoforms of human ApoE (NB110-60531) or mouse IgG isotype control to CNBr-activated Sepharose 4 fast flow beads for isolation of native ApoE lipoprotein particles. Antibody-coupled beads were washed with coupling buffer (0.1 M NaHCO_3_, 0.5 M NaCl, pH 8.3) and the unreacted groups were quenched with 0.1 M Tris-HCl, pH 8.0 with rocking for 2h at 20°C. Beads were further washed with 0.1 M Tris-HCl pH 8.0 with 0.5 M NaCl and finally the antibody-conjugated beads were washed with 1X PBS prior to immunoprecipitation of ApoE lipoprotein particles from ACM. Concentrated ACM was then incubated overnight at 4°C with antibody-conjugated beads with end-to-end mixing and mild agitation. The beads were washed in 0.5 M NaCl to remove nonspecific proteins and the native apoE lipoprotein particles were eluted using 3 M NaSCN for immunoblot analysis of native PAGE gel with human anti-ApoE (1:500 #ab52607).

#### Brain and spinal cord tissue processing

Four weeks after AAV injections, mice were euthanized with isoflurance and transcardially perfused with PBS followed by 4% PFA in PBS. Brains and spinal cords were dissected, embedded in 10% gelatin and postfixed in 4% PFA overnight at 4°C. Next day, gelatin embedded tissues were transferred to PBS with 0.05% sodium azide and stored at 4°C. Brains were then processed using vibratome (Leica VT1000S). Coronal sections (40 µm) of brain were processed using vibratome (Leica VT1000S) and stored in PBS with 0.05% sodium azide at 4°C. For spinal cord, we generated transverse or sagittal sections depending on the analysis. We used sagittal sections (40 µm) for scoring 5-HT raphespinal in lumbar caudal to injury site and CST tract labeling density at cervical region. Furthermore, a 10 mm block of spinal cord consisting of thoracic hemisection site (-5 mm rostral and +5 mm caudal) was excised and sectioned sagittal (40 µm) to access CST tract regeneration and severity of lesion. To analyze CST regeneration, sections underwent an immunohistochemistry protocol to enhance tdTomato signals for confocal microscopy. In brief, sections were incubated for 10 min in 0.3% H_2_O_2_ at room temperature, washed thrice in PBST (0.05% Triton-X) and blocked in 2% pre-filtered BSA with 5% normal donkey serum (Jackson ImmunoResearch: 017-000-121) and 5% normal goat serum (005-000-121) for 1 h at room temperature. Sections were then incubated overnight in Rb anti-mCherry (1:1000; ab167453) prepared in blocking solution mentioned above. Following day, the sections were washed 3 times in PBS-T and then incubated for 2 h in 568 D-anti-Rb (1:1000). For scoring raphespinal axons, sagittal sections were blocked as mentioned above and stained with Rb anti-5-HT serotonin 1A receptor (1:10000; Immunostar 24504). For scoring glial scar at injury site, the following antibodies were used monoclonal anti-GFAP 2.2B10 (1:1000 #13-0300); polyclonal anti-Iba1 (1:500 Wako); polyclonal anti-Laminin (1:1000 #PA1-167730). For brain coronal sections to analyze CST cell bodies in M1 cortex, following antibodies were used anti-Lrp8 (1:500 #4H3E6 MABN1872); human anti-ApoE (1:500 #ab52607); anti-NeuN (1:1000 A60 #ABN91); anti-ChAT (1:1000 AB143); anti-LRP5 (1:1000 36-5400); anti-LRP6 (1:2000 # PA5-101047); anti-Efnb2 (1:1000 # MA5-32740); anti-Tsc1 (1:1000 # 6935); anti-Synpo (1:500 # 163 002); anti-Homer (1:1000 # 160 023). After 3x washes in PBS-T, every other section was mounted on Superfrost Plus slide with VectorShield antifade medium (H-1200-10). At the end, fluorescent Nissl stain performed for one h at room temperature using NeuroTrace 435/455 Blue (1:1000, N21479). Sections were imaged birectional at 40x magnification using Leica SP8 Confocal at 1024-pixel resolution with 400 scan speed. Images were acquired titled and stitched using Leica SP8 post processing software for further morphometric analysis. The number of regenerating fibers in spinal cord sections were normalized to dorsal column CST bundle at level of cervical enlargement. Axonal counting and glial scar quantification was performed as previously described(*55*). Animals are chip-tagged and respective IDs were used for sampling and data collection and hence performed blind to genotypes.

#### Intensity measurements

For quantification of somatic Lrp8 signal intensity, we used ImageJ/FIJI(*58*). For each condition, we used *z*-projection of deconvolved stacks for two NeuN and Lrp8 channels. Using grey scale images, a nuclear mask was generated using NeuN pattern. The nuclear mask was then extracted and transferred to grey scale images of Lrp8 images and dilated twice to cover the Lrp8 signal distribution in each neuron. The difference between nuclear mask and twice dilated mask represents Lrp8 intensity in each soma. Instead of using cytoplasmic area by selecting the whole neuronal soma, here we use the nuclear mask and enlarge it several times to capture the cytoplasmic or membrane signal outside the nucleus. This approach has an advantage of preserving a uniform relationship between nuclear mask and cytoplasmic mask under most circumstances. Mean fluorescence intensity of Lrp8 were calculated for each section by dividing total fluorescence intensity by number of pixels measured for each soma. The intensity of synaptic proteins in neuropile was quantified using ImageJ/FIJI. After background subtraction, and the mean signal intensity across the whole image was measured for each protein.

#### Axon regeneration assay

Healthy uninjured mouse cortical neurons (7 DIV) and iPSC derived human neurons (45-50 DIV) were injured using a multipin scrapping tool and allowed to regenerate for another 7 d before fixation with 4% PFA in PBS. Axon regeneration in the scrape zone were visualized using anti-ßIII tubulin mAb (1:5000, G7121 Promega), phalloidin conjugates for actin staining (1;1000, A12379 and A12381, Invitrogen) and DAPI (1:5000, Bio-Rad). For scoring axon regeneration, images acquired at 10x magnification in an automated high-throughput imager (ImageXpress Micro confocal, Molecular Devices) under identical conditions for all experiments. Image thresholding and quantification were automated using ImageJ script angiotube formation algorithm to analyse axon regrowth(*59*). The entire dataset was manually blinded by the investigator before analysis.

#### Surface biotinylation and LRP8 recycling

Primary mouse cortical neurons were treated with purified secreted ApoE (12 µg/ml), 24 h after axotomy on day 7 of culture. On day 12, primary neurons were treated with Reelin (2 µg/ml) for 1 h before performing surface biotinylation assay (see timeline in Fig. 4C). After reelin treatment, the cells were washed with ice cold PBS and incubated in PBS containing sulfo-NHS-SS-Biotin reagent (1.0 µg/ml) for 30 min at 4°C. Rinsing the neurons again with ice cold PBS with 100 mM glycine helps to quench the excess biotin reagent. Neurons were then lysed using 10X RIPA lysis buffer (EMD Millipore 20-188) with EDTA-free protease and phosphatase inhibitors cocktail tablets (Roche 11873580001 and 04906837001) at 4°C for 20 min. Debris was removed by centrifugation for 15 min at 15,000 rpm at 4°C. Protein estimated using PierceTM BCA protein assay kit (#23225). 100 µg of total protein was incubated with 50 µl of NeutrAvidin agarose pellets at 4°C for 1 hr. Biotinylated surface proteins that are bound to be agarose pellets was washed thrice in washing buffer containing 500 mM NaCl, 15 mM Tris-HCl, 0.5% Triton X-100 at pH 8.0. Biotinylated surface proteins eluted from beads by boiling in 4X SDS sample loading buffer and analyzed on SDS-PAGE. The following antibodies were used for immunoblot: anti-ApoE (1:1000 ab52607), anti-LRP8 (1:500 4H3E6 MABN1872), anti-LRP5 (1:1000 D80F2 #5731), anti-LRP6 (1:500 C5C7 #2560), anti-Src (1:1000 36D10), anti-phospho-Src family (Tyr416, D49G4 1:1000). Immunoblots developed using Infrared dyes of 680 and 800 dyes in LI-COR Odyssey imaging systems. Single channel, high-resolution tiff images were converted to grey scale for densiometric analysis using Image Studio Lite image processing software.

#### Single nuclei 10x genomic sequencing and analysis

Mouse brain tissue collection and nuclei isolation for single nuclei RNA-seq were completed as previously described with slight modification(*60*). E2 and E4 mice were aged for 6 weeks and then underwent either a sham or dorsal thoracic over-hemisection surgical procedure. Eleven weeks post-surgery, E2 and E4 mice with and without SCI were sacrificed via rapid dissection. For each mouse, the motor cortex and corresponding sub-cortical regions from the right-brain hemisphere were micro-dissected, pooled, and immediately frozen on dry ice then stored at -80°C until nuclei isolation could be performed. Individual tissues were separately homogenized and then underwent density-based nuclei separation via centrifuging for one hour. Nuclei pellets were obtained, resuspended, and counted on a hemocytometer to achieve a concentration of 700-1200 nuclei/µl for generating single nuclei cDNA libraries.

Barcode-incorporated single-nucleus cDNA libraries were constructed using the Chromium Single Cell 3’ Reagents Kit v3 (10x Genomics) following the manufacturer’s guidelines. Sample cDNA libraries of all experimental groups were pooled and batched sequenced on an Illumina NovaSeq 5000 using single-indexed paired-end HiSeq sequencing. A sequencing depth of >400 million reads was achieved for all samples with an average read depth of 50,000 reads per nuclei. The resulting sequencing binary base call files (BCLs) were demultiplexed into FASTQ files. Sequenced samples were aligned to the mm10-2020-A *Mus musculus* reference genome using the Cell Ranger Count software (pipeline version 7.1, 10x Genomics), generating barcoded sparse matrices of nuclei-gene raw UMI counts.

Sample nuclei-gene count matrices were imported and combined into a single anndata object for quality control (QC) and downstream processing with Scanpy (version 1.9.3)(*61*), a scalable toolkit for Python-based gene expression analyses. Individual samples with total nuclei counts outside the interquartile range (IQR) of the data set were removed to improve the reliability of the results. Genes detected in less than 10 nuclei were discarded. Nuclei with over 5% of UMI counts mapped to mitochondria genes and nuclei with less than 50 genes or more than 5,000 genes detected were considered outliers and discarded. Raw nuclei-gene counts were retained as a separate layer within the annotated object for integrated clustering. For comparative gene expression analysis, the UMI counts were normalized to their library size by scaling to 10,000 transcripts per nucleus and then log-transformed. Across all experimental groups, 201,941 nuclei and 26,473 genes were retained post-QC processing and clustering.

In total, 20 samples were used for integrated clustering (E4, n=5; E2, n=4; E4-SCI, n=5; E2-SCI, n=6). Sample clustering was performed by first using Scanpy’s *highly_variable_genes* (HVGs) function, and the implementation of *‘flavor=seurat_v3’*, to identify shared HVGs between samples(*62, 63*). Briefly, this method computes a standardized normalized variance by which genes are ranked. Genes are then sorted by their variance and rank-matched between samples, with the top 2500 genes selected as HVGs based on the number of samples with shared rankings. Next, the scVI probabilistic model from the scVI-tools package was used for sample-based batch correction, incorporating covariate regression of total library size and mitochondrial transcripts per nucleus(*64*). The model was then trained to obtain the latent space representation of each nucleus.

A neighborhood graph of the batch-corrected matrix was constructed using Scanpy’s *neighbourhoods* function, with *‘n_neighbourhoods’* size of 15 and the scVI-generated latent space representation as input parameters. Integrated Leiden clustering of nuclei into cell-type subgroups was performed from the neighbourhood graph and dimensionally reduced in UMAP space for visualization. To enhance the clustering process, nuclei identified as doublets were excluded from the dataset, and the integrated Leiden clustering was repeated using a newly retrained scVI model. Doublets were identified based on clusters showing enrichment for marker genes of different cell types, suggesting mixed-cell expression profiles within the same cluster.

Clusters of nuclei enriched with specific marker genes were classified as corresponding primary cell types. To identify the top differentially expressed marker genes within individual Leiden clusters, the *rank_genes_groups* function in Scanpy was used with the Wilcoxon Ranked-Sum test. Genes with the highest ranks showing unique enrichment in a single cluster were considered marker genes. The cell type specificity of these markers was verified using previously published data(*60, 65, 66*).

Additionally, the *rank_genes_groups* function with the Wilcoxon Ranked-Sum test was employed for differential gene expression (DEG) analysis between experimental groups for each major identified cell type. Some sub-clusters, enriched with nuclei from a single sample, were excluded from all DEG analyses. Using the remaining clusters, only genes expressed in at least 10% of all nuclei post-ranking were considered differentially expressed. Significant DEGs were defined as those with an adjusted *p-value* (false discovery rate) of less than 0.05 and a Log_2_ fold-change greater than ±0.25, unless specified otherwise (Table S1).

Gene set enrichment analysis of DEGs from select cell types was conducted using Cytoscape (version 3.9.1)(*67*) with ClueGo (version 3.10.2)(*68*) extensions. DEG lists were analyzed in ClueGo to identify overrepresented functional and pathway terms using the Gene Ontology (GO) Biological Processes, Molecular Functions, and Reactome Pathways and Reactions databases. A two-sided hypergeometric test with BH-adjusted p-value of 0.005 and a 5-gene threshold was used for term enrichment. GO term fusion was applied to merge redundant and related biological themes with more than 50% gene similarity. Filtered term associations were visually organized into functionally related clustered network using the GeneMANIA-based clustering algorithm(*69, 70*). Selected GO-term associated gene subsets were further assessed for protein-protein interactions (PPI) using STRING (version 12)(*71, 72*), visualizing PPI networks with a confidence score of 0.3 for gene association strength.

#### Statistics

Statistical analysis was performed in Prism 10.3.1. All data presented here as mean ± SEM unless specified otherwise. Two-way ANOVA with repeated measures was performed SPSS for time course datasets of repeated measurements. Student’s two-tailed unpaired t-test and one-way ANOVA with Tukey’s multiple comparisons test were used for pairwise comparisons. Statistical significance was set as *p* < 0.05 for all analysis. The specific test used to calculate *p* value, number of replicates, F and Df are noted in Fig. legends.

## SUPPLEMENTARY FIGURES S1-S7

**Supplementary Fig. S1.**
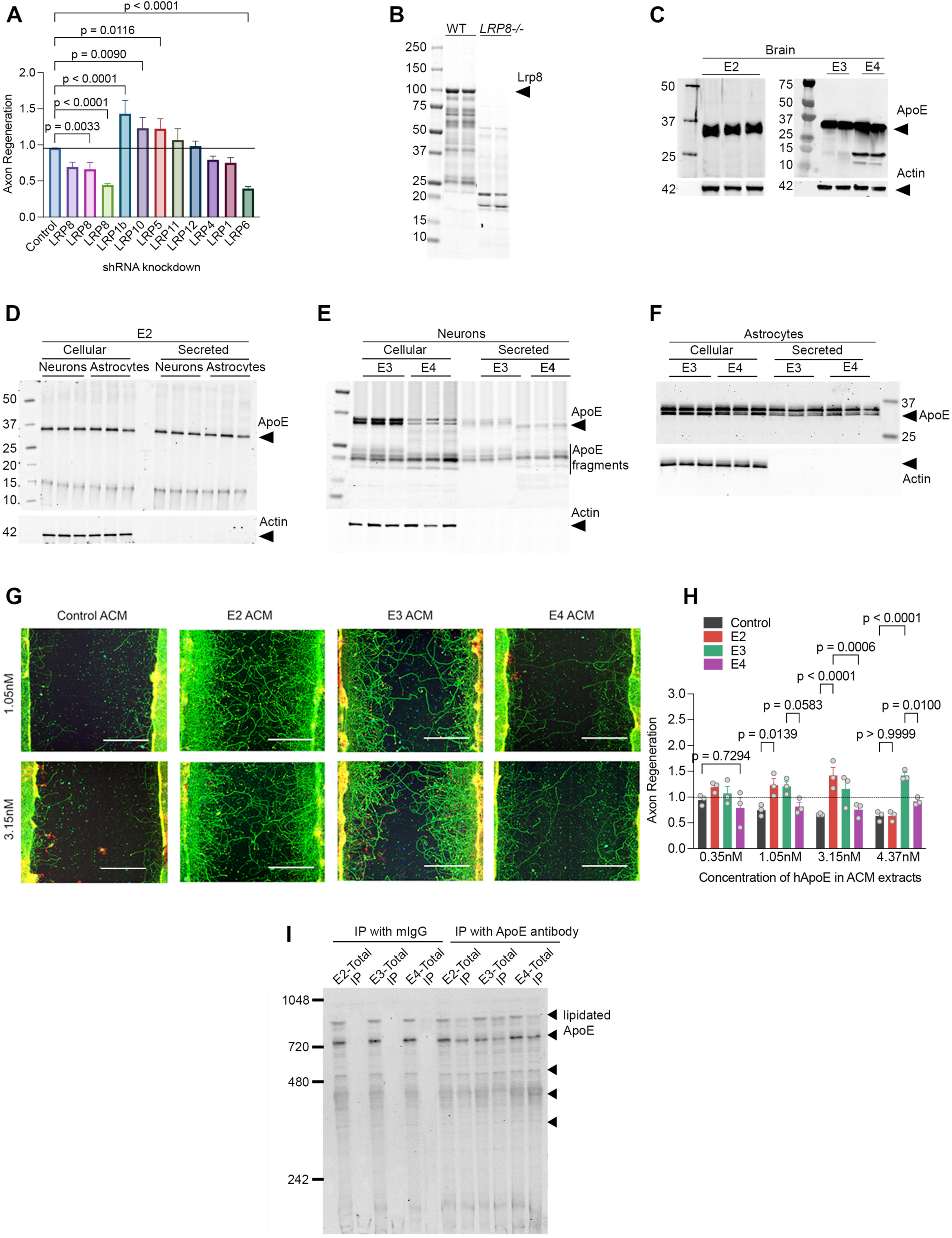
Role of ApoE-Lrp8 in axon regeneration. (A) Quantification of axon regeneration in WT cortical neurons cultured with shRNA-mediated knockdown for different members of LDL receptor-related protein (LRP) family. Data represents mean ± SEM from three biological replicates and analyzed with one-ANOVA with Dunnett’s multiple comparison test (F = 6.25, dF = 11). (B) Immunoblot of RIPA total protein extracts isolated from brain of WT and *Lrp8^-/-^* to access the specificity of LRP8 antibody. (C) Anti-ApoE immunoblot of RIPA extracted total protein isolated from mouse brain carrying the indicated human APOE-KI alleles. Anti-actin used as loading control. (D) Anti-ApoE immunoblot analysis of RIPA extracts from cellular and secreted fractions of cortical neuronal and astrocytes cultures from E2 mice. Anti-actin used as loading control. (E) Anti-ApoE immunoblot analysis of RIPA extracts from cellular and secreted fractions of cortical neuronal cultures from E3 and E4 mice. Anti-actin used as loading control. (F) Anti-ApoE immunoblot analysis of RIPA extracts from cellular and secreted fractions of cortical astrocytes cultures from E3 and E4 mice. Anti-actin used as loading control. (G) Photomicrographs of axon regeneration in cultured cortical neurons from EKO mice treated with astrocyte conditioned medium (ACM) containing 1.05 nM and 3.15 nM of hApoE derived from cortical astrocytic cultures from respective APOE-KI mice. Cultures stained at d15 for ßIII-tubulin (green) and phalloidin (red). Scale bar, 200 µm. (H) Quantification of axon regeneration from (G). Data shown as mean ± SEM from three biological replicates and analyzed by two-way ANOVA with Tukey’s multiple comparisons test with F (9, 32) = 4.5, p = 0.0007 for interaction effects, F (3. 32) = 18.45, p<0.0001 for different ACM effects and F (3, 32) = 0.86, p = 0.46 between different ACM concentrations. (I) Native PAGE analysis to detect ApoE lipoprotein profile after immunoprecipitation with mIgG and anti-hApoE using ACM extracts derived from astrocytic cultures of hAPOE-KI mice.

**Supplementary Fig. S2.**
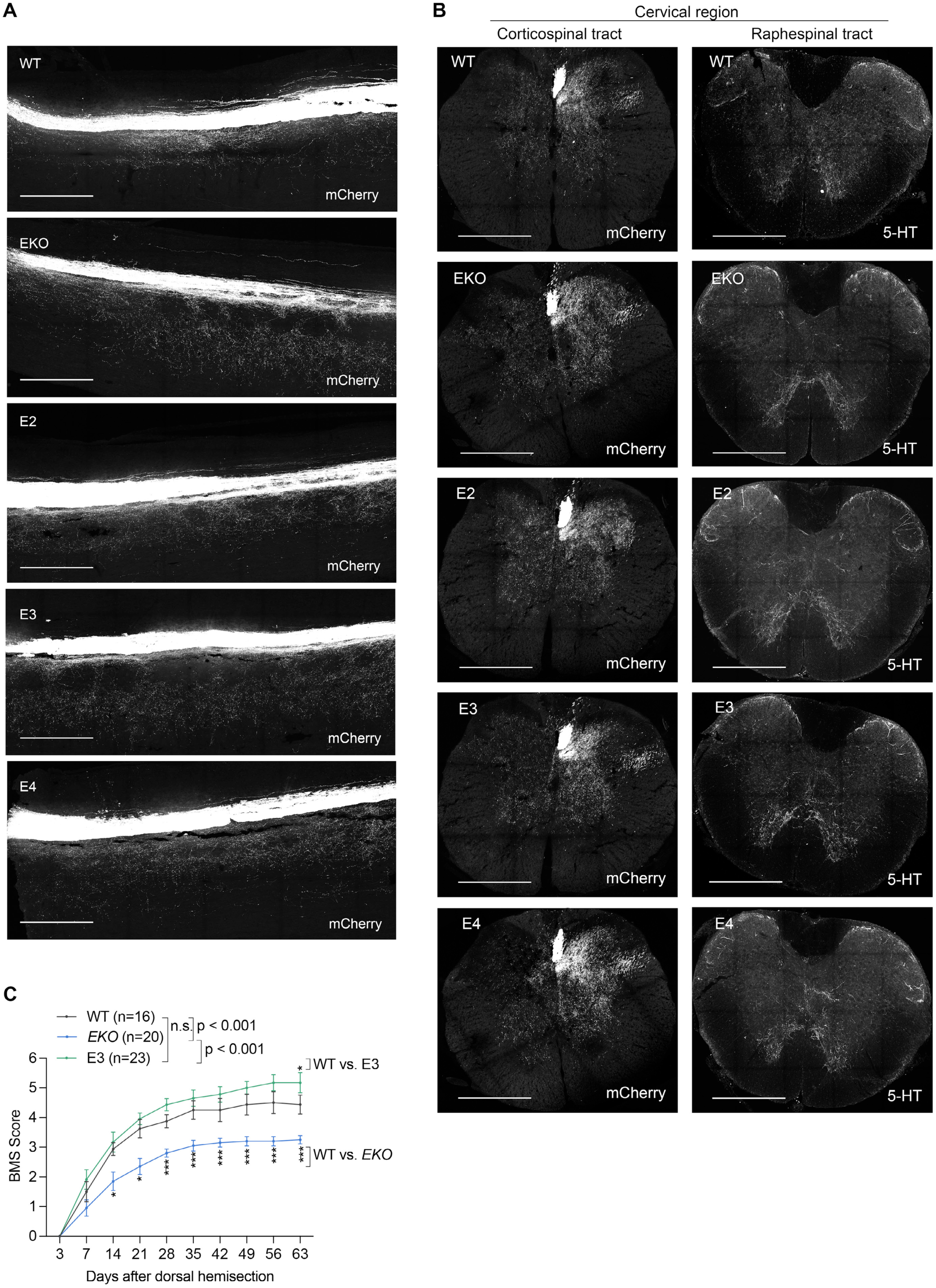
Development of corticospinal and raphespinal tracts in hAPOE-KI mice and its role in SCI recovery. (A) Sagittal photomicrographs of thoracic region of spinal cord. WT (*n* = 3), EKO (*n* = 4), E2 (*n* = 3), E3 (*n* = 3), E4 (*n* = 3) mice, injected into M1 cortex with AAV-based anatomical tracer to label CST axon bundle. Sections stained with anti-mCherry to access CST development. Dorsal is up and rostral is left. Scale bar, 200 µm. (B) Transverse photomicrographs of cervical region of spinal cord stained with anti-mCherry to access CST tract and with anti-5HT to access development of raphespinal tract in the experimental cohort described in Fig. 2A. Scale bar, 200 µm. (C) Open-field performance score for locomotion by BMS score of WT (*n* = 16), EKO (*n* = 20) and E3 (*n* = 23). Performance scores were recorded for each animal every consecutive week and data reported as mean ± SEM and analyzed using repeated measure ANOVA across time series followed by post hoc Tukey’s multiple comparisons test for genotypic effect at indicated time points. ***p<0.001, *p<0.05.

**Supplementary Fig. S3.**
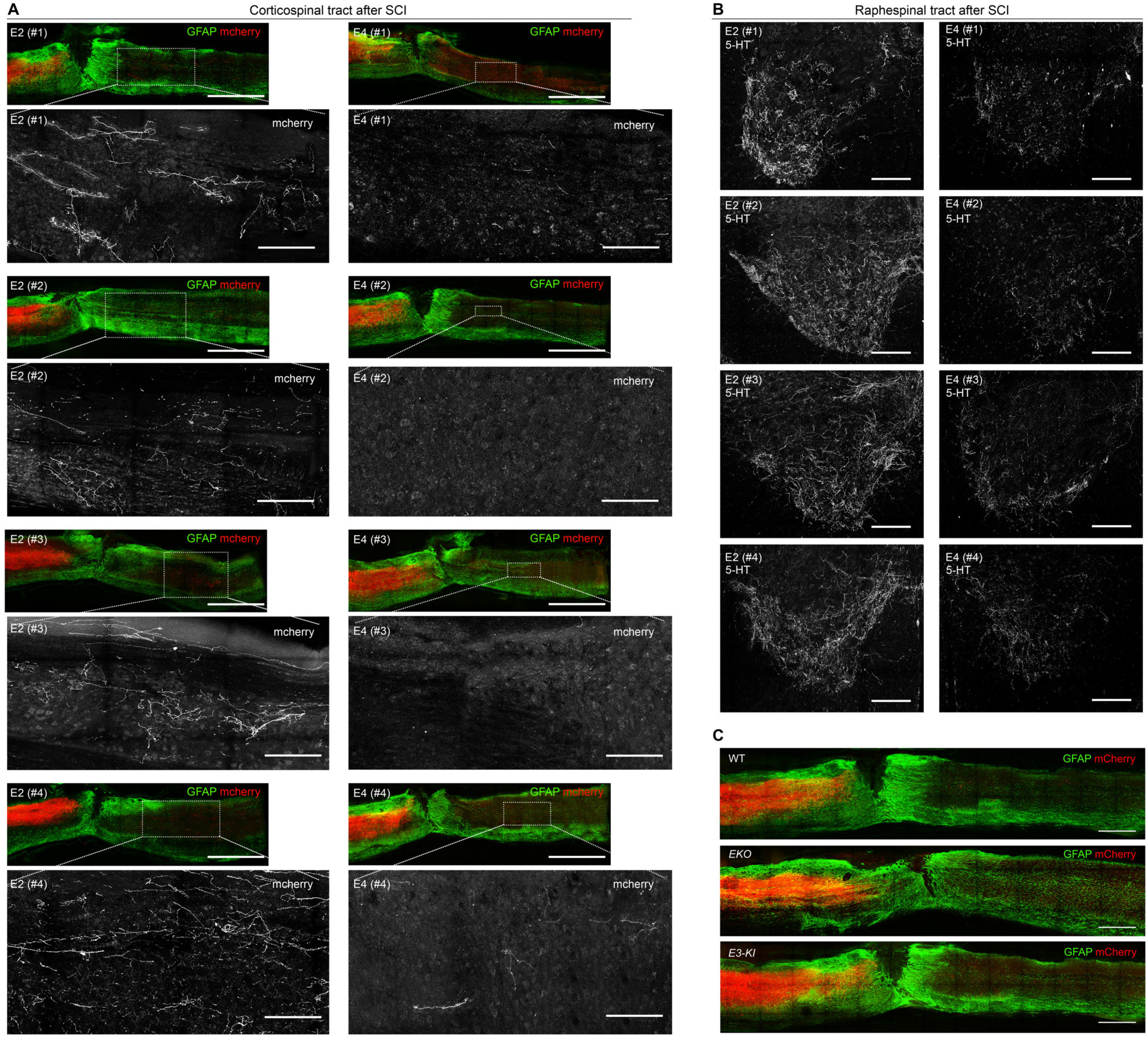
E2 mice exhibit pronounced reparative axon regrowth after SCI. (A) Sagittal low power photomicrographs of spinal cord around the lesion site in E2 and E4 mice. Sections were stained with anti-GFAP (green) and anti-mCherry (red). Dorsal is up and rostral is left. Scale bar, 500 µm. White outlined boxed areas in each image are captured at high resolution to visualize regenerating CST fibers caudal to lesion for red channel only. Scale bar, 100 µm. (B) Transverse section photomicrograph of ventral horn lumbar spinal cord of E2 and E4 mice. Sections stained with anti-5-HT to visualize raphespinal fibers. Scale bar, 100 µm. (C) Sagittal photomicrographs of spinal cord around the lesion site in WT (*n* = 12), *EKO* (*n* = 13) and E3 (*n* = 16) mice. Sections were stained with anti-GFAP (green) and anti-mCherry (red). Dorsal is up and rostral is left. Scale bar, 500 µm.

**Supplementary Fig. S4.**
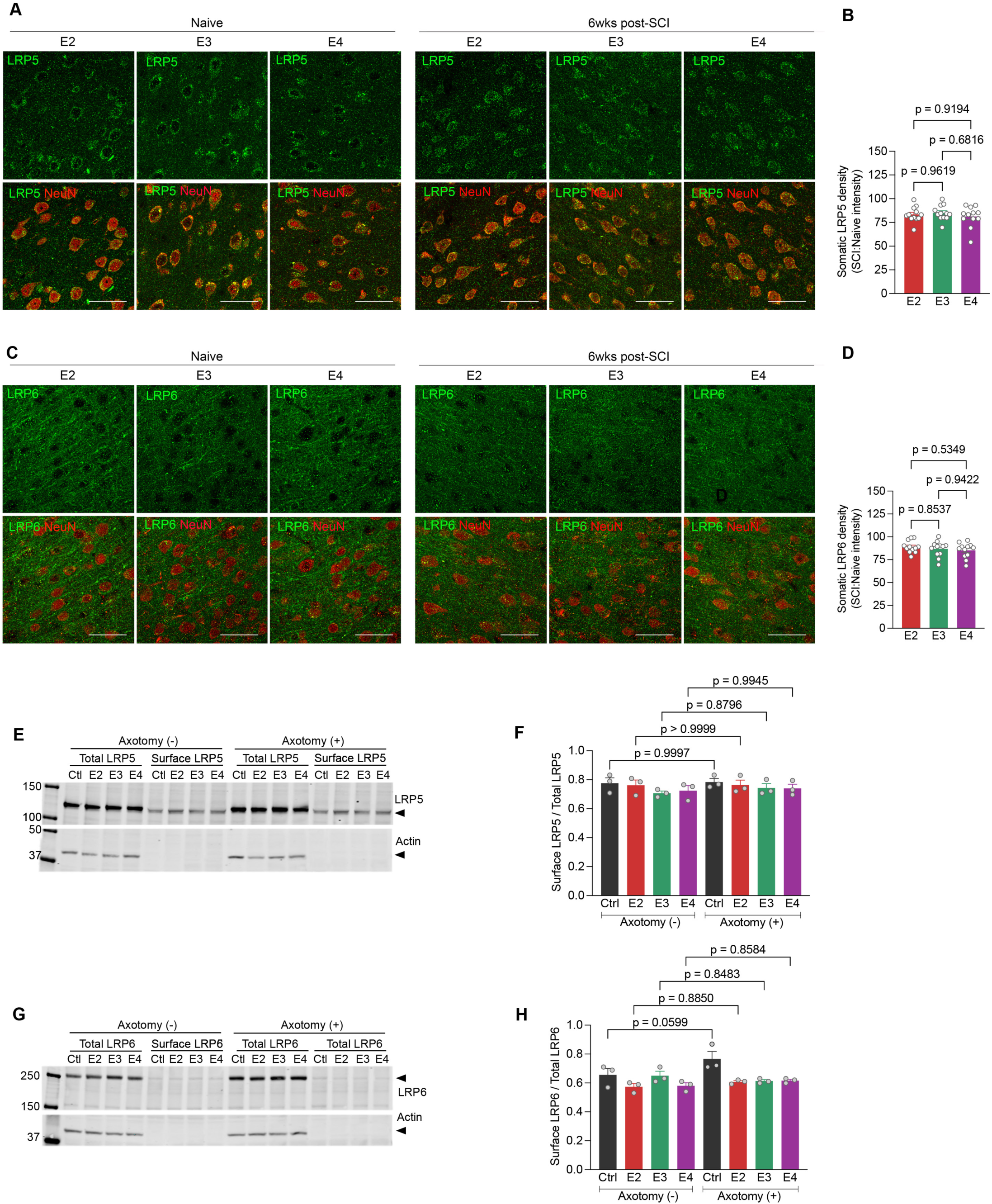
LRP5 and LRP6 receptors not required for ApoE dependent reparative axon growth. (A) Photomicrographs of L5 from M1 brain cortex stained for anti-LRP5 (green) and anti-NeuN (red) 42 days after SCI compared to age-matched naïve controls for respective genotypes. Scale bar, 200 µm. (B) Quantification of LRP5 levels in neuronal soma 42 days after SCI for E2, E3 and E4 mice (*n* = 12). Data presented as mean ± SEM and analyzed two-tailed unpaired t test (t = 0.54, dF = 22). (C) Photomicrographs of L5 from M1 brain cortex stained for anti-LRP6 (green) and anti-NeuN (red) 42 days after SCI compared to age-matched naïve controls for respective genotypes. Scale bar, 200 µm. (D) Quantification of LRP6 levels in neuronal soma 42 days after SCI for E2, E3, E4 mice (*n* = 12). Data presented as mean ± SEM and analyzed two-tailed unpaired t test (t = 1.29, dF = 22). (E) Immunoblot with total and biotinylated surface protein extracts from Fig. 4J to measure differential recycling of surface LRP5 with actin as loading control. (F) Densitometry analysis of immunoblots from three independent experiments described in Fig. 4J for surface LRP5 with excess reelin. Data presented as mean ± SEM and analyzed by one-way ANOVA with Tukey’s multiple comparisons test with F = 0.73, dF = 7. (G) Immunoblot with total and biotinylated surface protein extracts from (Fig. 4J) to measure differential recycling of surface LRP6 with actin as loading control. (H) Densitometry analysis of immunoblots from three independent experiments described in (Fig. 4J) for surface LRP6 with excess reelin. Data presented as mean ± SEM and analyzed by one-way ANOVA with Tukey’s multiple comparisons test with F = 4.53, dF = 7.

**Supplementary Fig. S5.**
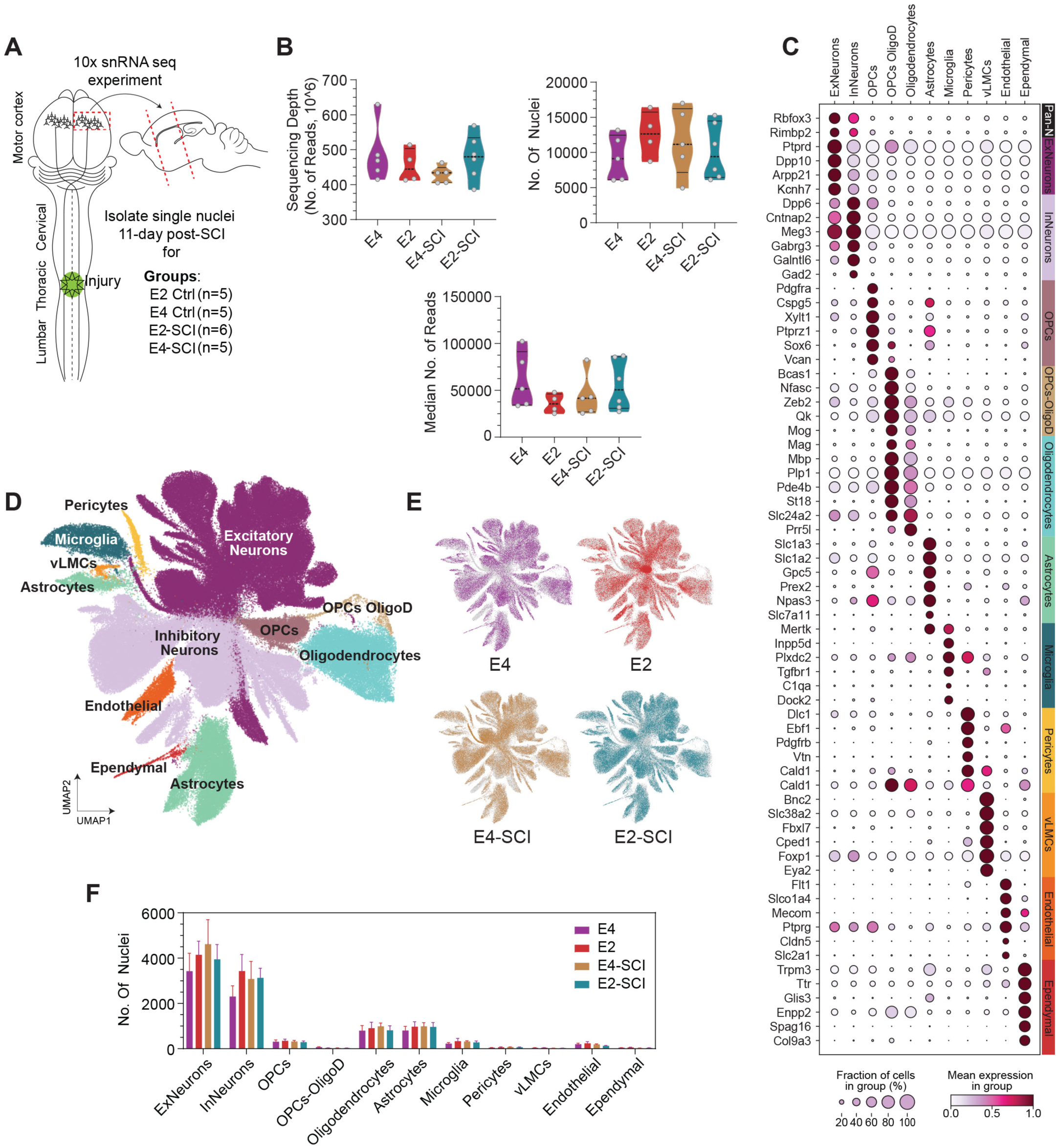
Integration of snRNA-seq datasets and classification of major cell types. (A) Schematic depicting the overview of experimental design, brain region used for single nuclei isolation and RNA sequencing across E2 and E4 experimental mice with pre- and post-injury. (B) The sequencing depth (left), number of total nuclei (right), and median number of reads (bottom) resulting from the single-nucleus RNA-sequencing (snRNAseq) of samples pooled by experimental groups. (C) UMAP of cell-type identified clusters. (D) Dot plot showing the percentage of nuclei and scaled mean expression of cell-type specific marker genes used to determine the UMAP clusters in C. (E) UMAP of collective sample gene expression profiles projections for each experimental group. (F) Comparative nuclei frequencies of clustered cell types of individual samples within each experimental group. Data is presented as the mean ± SEM and compared by a mixed model test with Tukey’s correction. No between-group statistical significance differences were detected.

**Supplementary Fig. S6.**
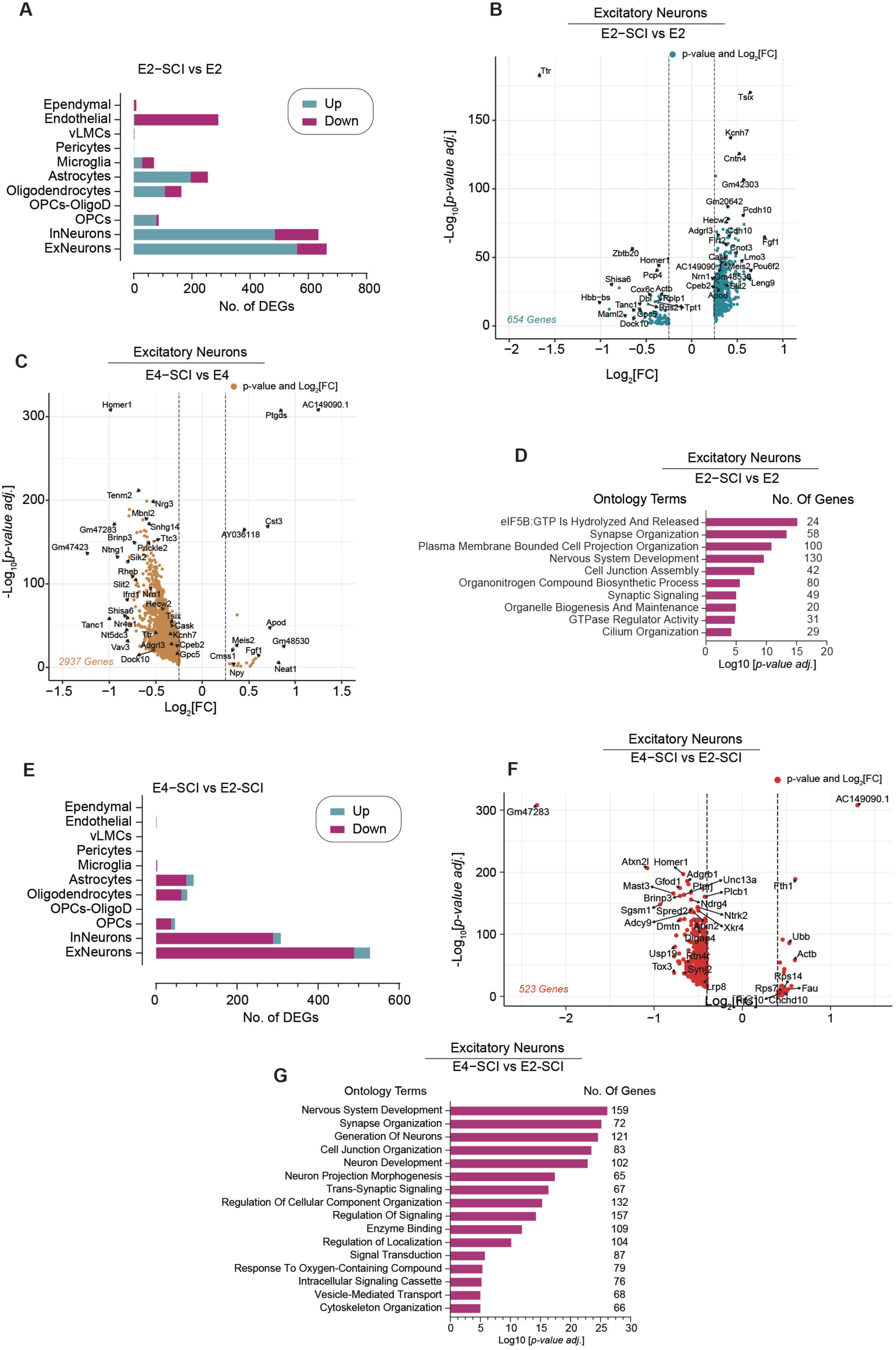
Cortical upregulation of SCI-induced growth-promoting transcriptomic signature is specific to the E2 allele. (A) The number of up and downregulated differentially expressed genes (DEGs) in neuronal and glial cell populations in E2 mice post-SCI. (B) Volcano plots representing the gene expression changes in the excitatory neuron populations of E2 mice post-SCI. (C) Volcano plots representing the gene expression changes in the excitatory neuron populations of E4 mice post-SCI. (D) Bar plot of the *p-*values, and corresponding number of term-associated genes, for the top leading terms from gene set pathway enrichment analysis of E2 excitatory neurons DEGs post-SCI shown in A. Complete list of enriched pathway terms can be found in Supplementary Table S3. (E) Number of up and down regulated DEGs in neuronal and glial cell populations in RAG category. (F) Volcano plot showing gene expression changes in excitatory neuron cells in RAG category. (G) Bar plot of the *p-*values, and corresponding number of term-associated genes, for the top leading terms from gene set pathway enrichment analysis of E2 excitatory neurons DEGs post-SCI shown in E. Complete list of enriched pathway terms can be found in Supplementary Table S4.

**Supplementary Fig. S7.**
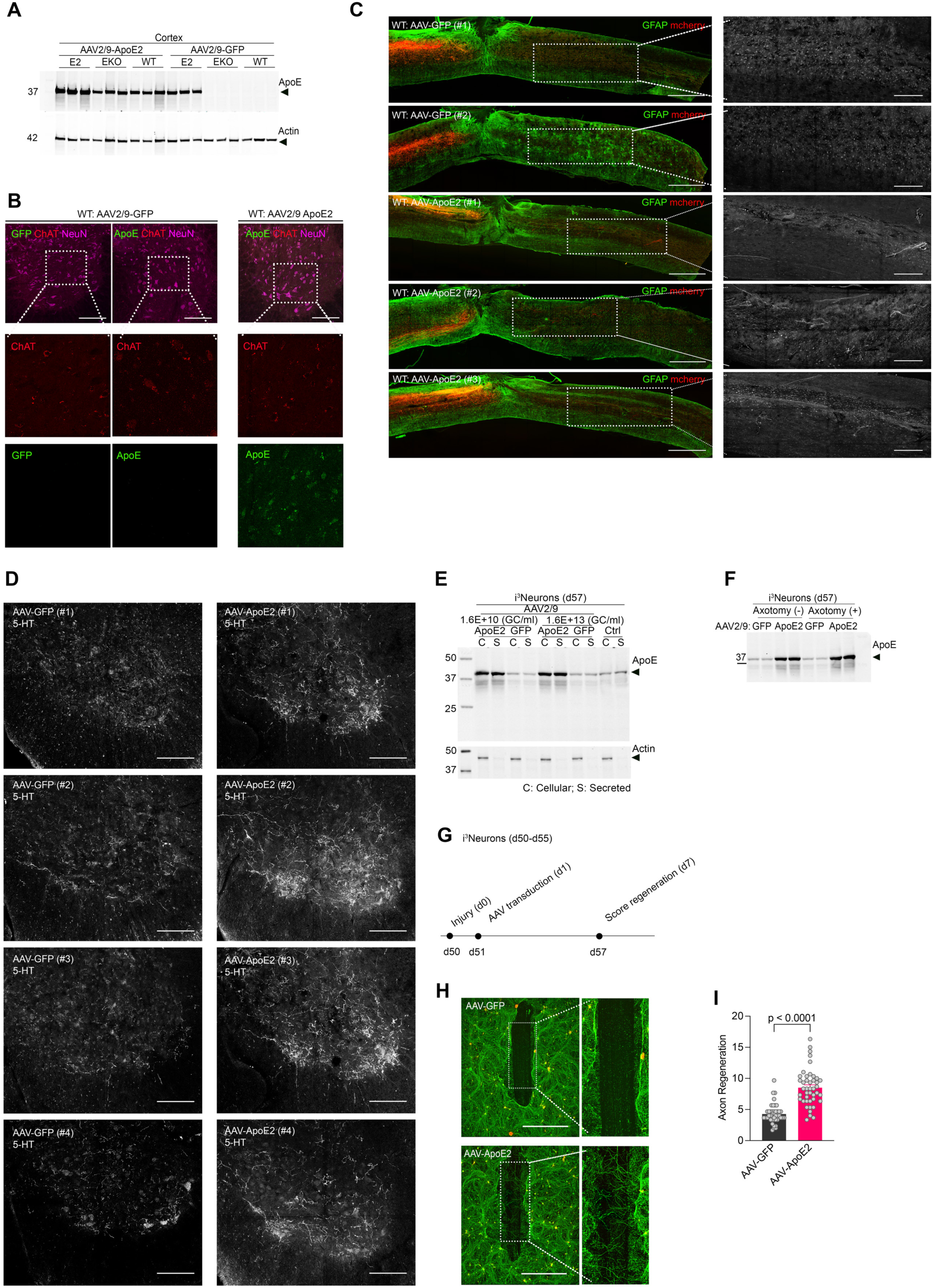
Expression profile analysis of AAV-mediated over expression of ApoE2 in cortex and i3N-derived human neurons. (A) Anti-ApoE immunoblot with total protein extracts from forebrain of WT, EKO and E2 mice with and without bilateral AAV2/9 injection at M1 cortex. Anti-actin used for loading control. (B) Transverse section photomicrograph of ventral horn lumbar spinal cord of WT mice at d77 (related to Fig. 7). Sections stained with anti-GFP, ApoE, ChAT and NeuN. Scale bar, 100 µm. (C) Sagittal low-power photomicrographs of spinal cord around the lesion site in AAV injected WT mice at d77 after SCI. Sections were stained with anti-GFAP (green) and anti-mCherry (red). Dorsal is up and rostral is left. Scale bar, 500 µm. White outlined boxed areas in each image are captured at high-resolution to visualize regenerating CST fibers caudal to lesion for red channel only. Scale bar 100 μm. (D) Transverse section photomicrograph of ventral horn lumbar spinal cord of WT mice expressing GFP and ApoE2. Sections stained with anti-5-HT. Scale bar, 100 µm. (E) Anti-ApoE immunoblot profile with cellular and secreted extracts of AAV2/9 transduced i3N neurons at d57. GFP expression is used as control to access levels of expressed ApoE2 compared to endogenous ApoE. Anti-actin used as loading control. (F) Anti-ApoE immunoblot profile with cellular extracts of AAV-GFP and ApoE2 transduced i3N neurons with and without axotomy. (G) Schematic to evaluate therapeutic benefit of ApoE2 in i^3^N-derived human neurons. (H) Photomicrograph showing regeneration zone of axotomized i^3^Neurons stained for ßIII-tubulin (green) and phalloidin (red). White outline in the axon regeneration zone is enlarged. (I) Quantification of axon regeneration index for i^3^N neurons expressing AAV-GFP and AAV-ApoE2 expression. Datapoint refers to each well from three independent replicates performed with different i^3^N-iPSC clones. *p* values calculated by two-tailed unpaired t-test with Welch’s correction (t = 8.35, dF = 64).

